# Revealing interactions between glutathione peroxidase 4 and phosphoinositides

**DOI:** 10.64898/2026.06.17.732910

**Authors:** Sara H. Walters, ByungUk Park, Rachel L. Signorelli, Courtney L. Labrecque, Faik N. Musayev, Reid C. Van Lehn, Brian Fuglestad

## Abstract

Glutathione peroxidase 4 (GPx4) is the primary enzyme that reduces lipid hydroperoxides to prevent membrane oxidative damage and protect against ferroptosis. GPx4 is known to engage with lipid headgroups through electrostatic interactions, positioning the substrate for reduction. This work reveals and characterizes binding of highly anionic phosphoinositides (PIP lipids) by GPx4. PIPs are vital lipids in human cells and are central to many signaling processes, particularly in cytosolic-facing membranes. Lipid overlay assays confirm interactions between GPx4 and phosphorylated PIPs, comparable to known anionic lipid binders. Protein NMR describes the interaction between GPx4 and PIPs within micelles. The greatest resonance shifting occurs with tris-phosphorylated PIP, suggesting that higher anionic charge leads to greater binding, a known driver of GPx4 substrate recognition. Preferred anionic interactions were also confirmed with titration and crystallographic structure analysis of *D-myo*-inositol 1,3,4,5-tetrakisphosphate (IP_4_). A headgroup-binding site on GPx4 is revealed to be proximal to the cationic membrane interaction site. In conjunction with molecular simulations, these results show that PIP lipid interactions allow full GPx4 membrane engagement and position the headgroup to allow the lipid tail to interact with the catalytic site. Understanding whether GPx4 preferentially interacts with PIPs will allow better understanding of the protective function of this essential enzyme and a mechanism that may protect essential lipid signaling pathways from oxidative damage.

**Significance:** This study allows a deeper structural and mechanistic understanding of GPx4, the primary enzyme that reduces lipid hydroperoxides and prevents ferroptosis. Gaining an understanding of phosphoinositide binding to GPx4 reveals a mechanism for potential preservation of these important signaling molecules and for ferroptosis protection. Observation of a specific binding site for headgroup engagement reveals a plausible lipid interaction mode and functional mechanism of this important cytoprotective enzyme.

## Introduction

Glutathione peroxidase 4 (GPx4) is a seleno-enzyme with isoforms localized to the cytosol, nucleus, and mitochondria of human cells.^1–3^ The cytosolic isoform of GPx4 is the primary enzyme responsible for reducing lipid hydroperoxides to prevent ferroptosis, a form of regulated cell death due to oxidative damage of lipids.^4,5^ A main driver of ferroptosis is the polyunsaturated fatty acyl (PUFA) tails of lipids, which are particularly susceptible to oxidative damage.^6,7^ Lipid types implicated in ferroptosis include phosphatidylcholine (PC), phosphatidylinositol (PI), and phosphatidylethanolamine (PE), especially when arachidonic acid or other PUFAs are incorporated or when modified to di-PUFA lipids.^6,8,9^ GPx4 eliminates hydroperoxides and limits the potential damage that can be caused from the iron-dependent propagation of lipid reactive oxygen species (ROS), maintaining membrane and cell homeostasis.^10^ GPx4 has recently become a highly sought after drug target due to its central role in ferroptosis regulation in a range of cancer cells, including mesenchymal-derived cancers, clear cell carcinomas, and persister cells from which a variety of drug-resistance mechanisms emerge.^11–13^ GPx4 inhibition and ferroptosis induction is lethal to many ferroptosis-sensitive cancers, indicating a novel therapeutic strategy.^14–16^ Mutations of GPx4 are medically relevant as well with the R152H mutation linked to Sedaghatian-type spondylometaphyseal dysplasia (SSMD) through reduced enzymatic activity of the protein due to a disruption in the active site, possible destabilization of the protein fold, and a change in its ability to anchor to membranes.^17–19^ Despite the biological and biomedical importance of GPx4, high-resolution experimental descriptions of substrate engagement are lacking. Further investigation of lipid processing and functional interactions by GPx4 is necessary to fully understand its molecular function and maximize its therapeutic potential.

GPx4 is a peripheral membrane protein (PMP); a water-soluble protein that interacts with membranes to perform its function.^20^ A combination of non-specific electrostatic and hydrophobic interactions are implicated in membrane binding by GPx4.^18,20–22^ Modeling and NMR studies have identified a cationic region adjacent to the active site, which is key to membrane engagement and lipid substrate binding.^20,21,23^ Anionic lipid headgroups bind to the cationic site (**Figure 1**, blue), which positions the hydroperoxidated lipid tail towards the catalytic site (**Figure 1**, red), allowing reduction. A number of basic residues within the cationic site are implicated in membrane binding or lipid substrate engagement, including K125, K127, K140, K151, and R152 (**Figure 1**). Reduced electrostatic interactions are implicated in a diminished ability of the SSMD R152H mutant GPx4 to engage with CL, confirming this site to be vital in lipid interactions and functionality.^18^ The highly anionic cardiolipin (CL) was found to be a high affinity binder compared to other phospholipids with lower anionic charge.^20,21^ Among the most negatively charged membrane lipids are the phosphatidylinositol phosphates (PIPs or phosphoinositides), which may contain up to three phosphorylations on the headgroup, in addition to the phosphodiester linking the inositol headgroup to the acylated glycerol. PIPs have a variety of possible phosphorylation patterns on the inositol headgroup in positions 3, 4, and/or 5. The negative charge of the headgroup increases with the addition of phosphorylated substituents, making them a potential target of the cationic site of GPx4. A major function of PIP lipids in cells is to interact with proteins.^24,25^ PIP lipids serve as signaling molecules for a variety of protein mediated cellular processes in the membrane, often by recruiting and regulating proteins at the cytosol-membrane interface.^24,26,27^ While PI was found to be among the higher affinity binders of GPx4,^21^ to our knowledge, the interaction of phosphorylated PIs has not been investigated. PIPs are often difficult class to study with both *in vivo* methods *in vitro* methods due to PIP lipids having varied, extremely low concentrations in cell membranes.^28^ Due to the heterogeneity and low abundance of these lipids, their protein interactions are often difficult to observe.^29^

**Figure 1.**
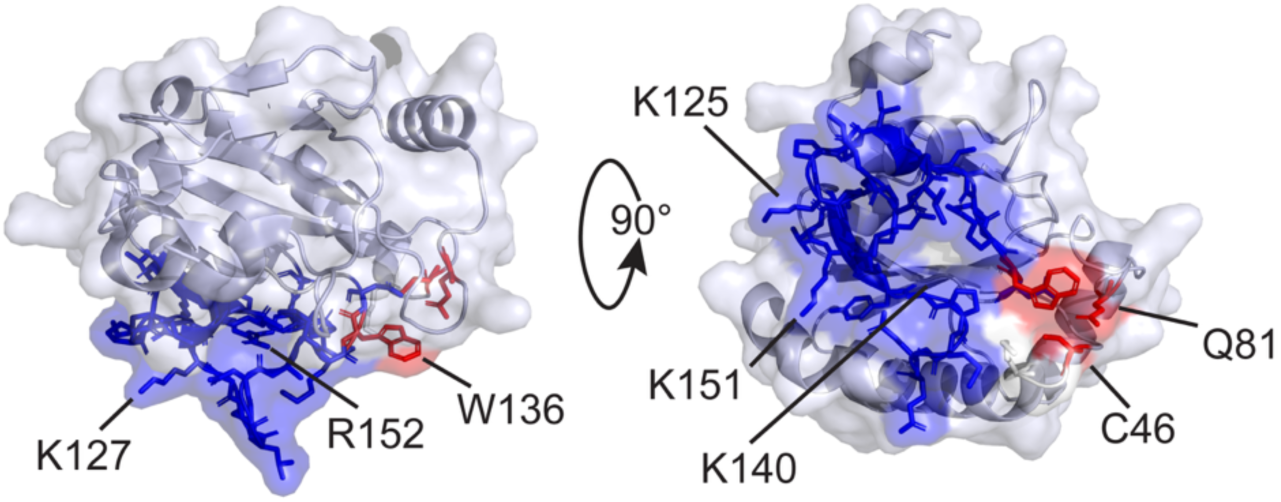
Depiction of the sites relevant to function in GPx4 (PDB: 2OBI).^21,22,26^ Blue indicates the cationic site which is responsible for membrane binding and lipid headgroup engagement and consists of residues 122-140, and 150-158, with important cationic residues labeled. Red indicates active, catalytic triad residues C46, Q81, and W136, which are labeled.

PIP lipids are most enriched in the inner leaflet of the plasma membrane, making them easily accessible to cytosolic GPx4.^28^ PIP species are generated from kinases and phosphatases which add or remove phosphates from the inositol headgroup.^30^ There has not yet been a link between PIP lipid hydroperoxidation and GPx4 reduction, possibly due to experimental constrains and difficulty in detection of PIPs.^31^ However, there are some contexts which provide a biological foundation for this study. Certain biological processes expose PIP lipids to oxidative damage, highlighting the need for GPx4’s protective function. For example, lipoxygenases (LOX) hydroperoxidate lipids for signaling or metabolic purposes, making them an antithesis to GPx4, with both proteins being important for cell homeostasis.^4,32^ LOXs (including 5-LOX, p12-LOX, and 15-LOX-1) all contribute to initiating cell death through ferroptosis.^33^ When GPx4 is active, it serves to counteract lipid hydroperoxidation formation from LOX enzymes, preventing propagation of oxidative damage from these LOX products.^33^ PIPs are the preferred substrates of some LOXs, specifically 15-LOXs, making potential GPx4 reduction of PIP-hydroperoxides important for protection from ferroptosis.^32^ Another key biological event where PIP hydroperoxidation may be expected is after phagocytosis where microorganisms are removed from the cell in a phagosome.^34^ PIP lipids are not only highly enriched in the phagosomal membrane but assist in initiating phagocytic reactive oxygen species (ROS) production, an event which may be expected to lead to oxidative damage of PIPs.^35^ In the plasma membrane, PIPs are highly enriched compared to PI, in contrast to other cell membranes where PI is the most prevalent isoform.^31,36^ Despite the overall low percentage of phosphorylated PIPs among all cell membranes, they account for 1-5% in the plasma membrane and play a significant role in protein binding.^29,37^ Due to their important role in signaling, reduction of PIP lipid hydroperoxides by GPx4 may serve to preserve intricate signaling networks. Additionally, phosphatidylinositol 3-kinase (PI3K), which is responsible for synthesizing phosphatidylinositol 3,4,5-trisphosphate (PI(3,4,5)P_3_), is known to be upregulated in a wide range of cancers.^38,39^ Therefore, PI(3,4,5)P_3_ may be particularly important in ferroptosis in cancer cells.^40^ With these considerations, understanding the relationship between PIPs and GPx4 is critical to a more complete understanding of the biology and therapeutic opportunities of ferroptosis.

Results here reveal interactions between GPx4 and PIPs, particularly PI(3,4,5)P_3_. The interactions are first verified through lipid overlay assays (LOAs), qualitatively showing that all phosphorylated PIP lipids interact with similar strength as lipids including CL, phosphatidylglycerol (PG), phosphatic acid (PA), and phosphatidylserine (PS), all previously implicated as preferred substrates of GPx4.^21^ Protein NMR data confirm GPx4 interactions with mono-, bis-, and tris-phosphorylated PIPs in the context of n-dodecylphosphocholine (DPC) micelles, with PI(3,4,5)P_3_ inducing the largest degree of shifting. Headgroup binding and recognition with inositol phosphates and other lipid headgroup analogs were tested to determine affinity differences based on headgroup and charge, resulting in the highest binding affinity with the PI(3,4,5)P_3_ headgroup analog, IP_4_ (1,3,4,5). This led to the observation of a headgroup binding site adjacent to the membrane interaction surface on GPx4. The headgroup interaction is confirmed by the ligand-bound crystallographic structure of GPx4 with IP_4_ (1,3,4,5). Molecular dynamics simulations give insight into how GPx4 functions on the membrane and as a hydroperoxidase. The PIP-GPx4 interaction was simulated with and without membrane models and confirmed the specific headgroup interactions identified by protein NMR and crystallography while additionally observing the interaction of the peroxidized lipid tail with the catalytic site of GPx4. Together, these results highlight a potential substrate preference for PIPs by GPx4 that have broad implications in ferroptosis prevention and protection of lipid signaling networks by lipid-hydroperoxide reduction.

## Results

We sought to understand whether GPx4 interacts with PIP lipids similarly to lipids such as CL or other known anionic binders.^18,21^ Interactions between PIP lipids and GPx4 were observed in two LOAs which report qualitative, comparative interactions of proteins against a variety of lipids. In this study, we utilized the U46C mutant of the cytosolic isoform of GPx4 throughout (referred to here as GPx4), where the native selenocysteine (position 46) is mutated to a cysteine (Figure 1). ^41,42^ Using the U46C mutant facilitates recombinant expression while retaining the wild-type structure, membrane and lipid binding, and some catalytic activity.^20,43,44^ The first LOA was performed using a Membrane Lipid Strip (Echelon Biosciences, **Figure 2A**). The anionic lipids CL, phosphatidylglycerol (PG), phosphatidylserine (PS), and phosphatidic acid (PA) were shown to have a relatively high affinity to GPx4 (**Figure 2A**). Sulfatide and PI also showed some binding with GPx4, but to a lower degree. The Membrane Lipid Strip also confirmed that the affinity of GPx4 to PE and PC was lower than the other tested lipids with no observable LOA signal.^21^ The phosphorylated PIPs, phosphatidylinositol 4-phosphate (PI(4)P), phosphatidylinositol 4,5-bisphosphate (PI(4,5)P_2_), and PI(3,4,5)P_3_ all showed high intensity comparable to CL, suggesting that PIP lipids may be preferred substrates of GPx4. A LOA assay using a PIP Strip (Echelon Biosciences) assayed a wider variety of immobilized PIP isoforms that may be present in cells and cell membranes, each with different patterns of phosphorylation. All PIP lipids displayed a qualitatively high comparable affinity to GPx4 (**Figure 2B**). However, this result must be interpreted with care since the immobilized, homogenous PIP lipids do not necessarily reflect a physiological, heterogeneous membrane. Nevertheless, LOA results suggest that GPx4 may interact with PIP lipids to a similar degree as other known anionic lipid substrates.

**Figure 2.**
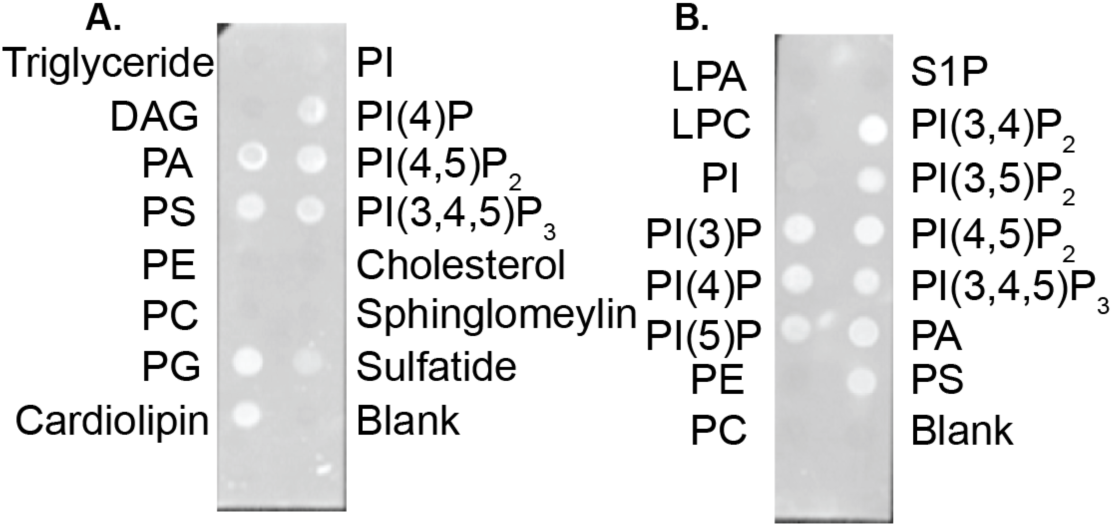
Lipid overlay assays confirm the binding of PIP lipids to GPx4. **A.** Membrane Lipid Strip (Echelon Biosciences) uses 100 pmol lipid/spot and confirms the binding of the PIP lipids along with PA, PS, PG, and cardiolipin (a known binder to GPx4). **B.** The PIP Strips (Echelon Biosciences) uses 100 pmol/spot of lipid and qualitatively show comparable binding affinity of GPx4 towards all phosphorylated PI lipids as well as PA and PS, which were shown on the Membrane Lipid Strip. Abbreviations: Diacylglycerol (DAG); Phosphatidic Acid (PA); Phosphatidylserine (PS); Phosphatidylethanolamine (PE); Phosphatidylcholine (PC); Phosphatidylglycerol (PG); Phosphatidylinositol (PI); Lysophosphatidic Acid (LPA); Lysophosphocholine (LPC); Sphingosine-1-phosphate (S1P); Phosphatidylinositol (3)-phosphate (PI(3)P); Phosphatidylinositol (4)-phosphate (PI(4)P); Phosphatidylinositol (5)-phosphate (PI(5)P); Phosphatidylinositol (3,4)-bisphosphate (PI(3,4)P_2_); Phosphatidylinositol (3,5)-bisphosphate (PI(3,5)P_2_); Phosphatidylinositol (4,5)-bisphosphate (PI(4,5)P_2_); Phosphatidylinositol (3,4,5)-trisphosphate (PI(3,4,5)P_3_).

To confirm the increased preference for PIP lipids, we titrated a water-soluble version of trisphosphorylated PIP, PI(3,4,5)P_3_ diC4. Results from a ^1^H-^15^N Heteronuclear Single Quantum Coherence (HSQC) NMR titration experiment (**Supplementary Fig. S1**) show shifting of resonances, indicating that GPx4 does interact with the water-soluble PIP analog.^20^ Estimating the apparent K_d_, based the high shifting residues from the chemical shift perturbations (CSPs), of the water-soluble PI(3,4,5)P_3_ diC4 is 4.6 ± 2.1 mM. Analogs of PS, PG, and PE headgroups were also added to aqueous GPx4 at 2 mM, but there were no visible or significant CSPs indicating a lack of binding, similarly to the previously observed PC water soluble headgroup (**Supplementary Figs. S2-4**).^20^ Though weak, this result confirms a preferential PIP-GPx4 interaction. Together, these results indicate that GPx4 binds to PIP lipids at least similarly to other known anionic lipid substrates.

To further characterize the interaction of PIPs with GPx4 in a more membrane-relevant context, we tested binding in micelles composed of n-dodecylphosphocholine (DPC) as the membrane model.^20,45^ Previous results showed that GPx4 retains its overall structure when bound to DPC micelles, demonstrated by minimal resonance shifting in regions that are not in contact with the micelle.^20^ In addition, function is retained in DPC micelles, as evidenced by the ability of GPx4 to reduce fatty acid hydroperoxides within the DPC micelle. These results demonstrate that DPC micelles are an adequate model for membrane-bound GPx4.^20^ Before PIP binding experiments, NMR resonance assignments of the U46C mutant of GPx4 in DPC micelles were mapped based on the published U46G-GPx4 assignments in a DPC micelle.^20^ Collection of 3D HNCA and HN(CO)CA experiments^46^ of DPC-bound GPx4 allowed confirmation and extension of mapped assignments and resulted in 75% of resonances assigned excluding prolines (**Supplementary Fig. S5**). The majority of unassigned residues are in the membrane-binding region and had reduced 3D NMR signal due to line-broadening, which are potentially related to functional dynamics.^47,48^ Observed resonance shifting confirms the membrane and lipid interactions; however, some residues that may respond to PIP binding are likely not observed. Nevertheless, a number of resonances within the membrane surface remain, allowing observation of comparative binding strength. NMR was used to observe GPx4 bound to DPC micelles and with and without the addition of 4:1 (mol/mol) lipid:protein for PI(3)P diC16, PI(3,4)P_2_ diC16, or PI(3,4,5)P_3_ diC16 to GPx4. Interpretable ^1^H-^15^N HSQCs of each micelle were successfully collected (**Figure 3A-C**). The NMR results reflected the anticipated shifting of observable GPx4 residues within the membrane interface. PI(3)P had the smallest degree of CSPs, PI(3,4)P_2_ had a higher degree of CSPs, while PI(3,4,5)P_3_ had the largest observed CSPs, suggesting that a greater number of phosphorylations result in stronger interactions with GPx4 in the context of a DPC micelle (**Figure 3D-F**). The addition of PI(3)P diC16 to the DPC micelle resulted in a 20% trimmed mean plus 1σ CSPs of 0.013 ppm (**Figure 3D**). The CSPs then increased with the addition of PI(3,4)P_2_ diC16 to the DPC micelle (**Figure 3E**) with a 20% trimmed mean plus 1σ of 0.020 ppm. The addition of PI(3,4,5)P_3_ diC16 to the DPC micelle resulted in C10, A11, S85, K90, A113, L116, K118, K121, K125, F138, L142, R152, G154, I162, and K164, all shifting greater than the 20% trimmed mean plus 1σ of 0.028 ppm (**Figure 3F**). Using the trimmed mean plus 1σ of the PI(3,4,5)P_3_, the PI(3)P residues above that threshold include only C10 and A11 and the PI(3,4)P_2_ residues above that threshold include C10, A11, S44, L116, K125, L142, R152, and I162. The residues in the region between A113-K121 border the cationic site, while residues between K121-N137 are a part of the cationic site.^20^ Residues between V150 and E158 are also a part of the cationic site and have a strong influence on membrane binding.^19,20^ The CSP results show that PIP lipids interact within the anticipated binding site, confirming the cationic region having an increased affinity to PIP lipids due to their high negative charge. 500 μM 12:0 PE added to the DPC micelle was used as a control displaying minimal CSPs (**Supplementary Fig. S6**), confirming the PIP interactions are exclusively from the PIP lipids and not simply due to an increased concentration of phospholipids. Likewise, a series of phospholipids within DPC micelles were tested by protein NMR. 1,2-dipalmitoyl-sn-glycero-3-phosphoserine (16:0 PS), 1,2-dilauroyl-sn-glycero-3-phosphate (12:0 PA), 1,2-dilauroyl-sn-glycero-3-phospho-(1’-rac-glycerol) (12:0 PG) displayed minimal spectral shifting at 500 μM when incorporated into DPC micelles (**Supplementary Figs. S7-9**). While these experiments provide evidence for preferential PIP binding within the GPx4-membrane interface, the interaction surface is only partially characterized due to incomplete assignments in this region of the protein.

**Figure 3.**
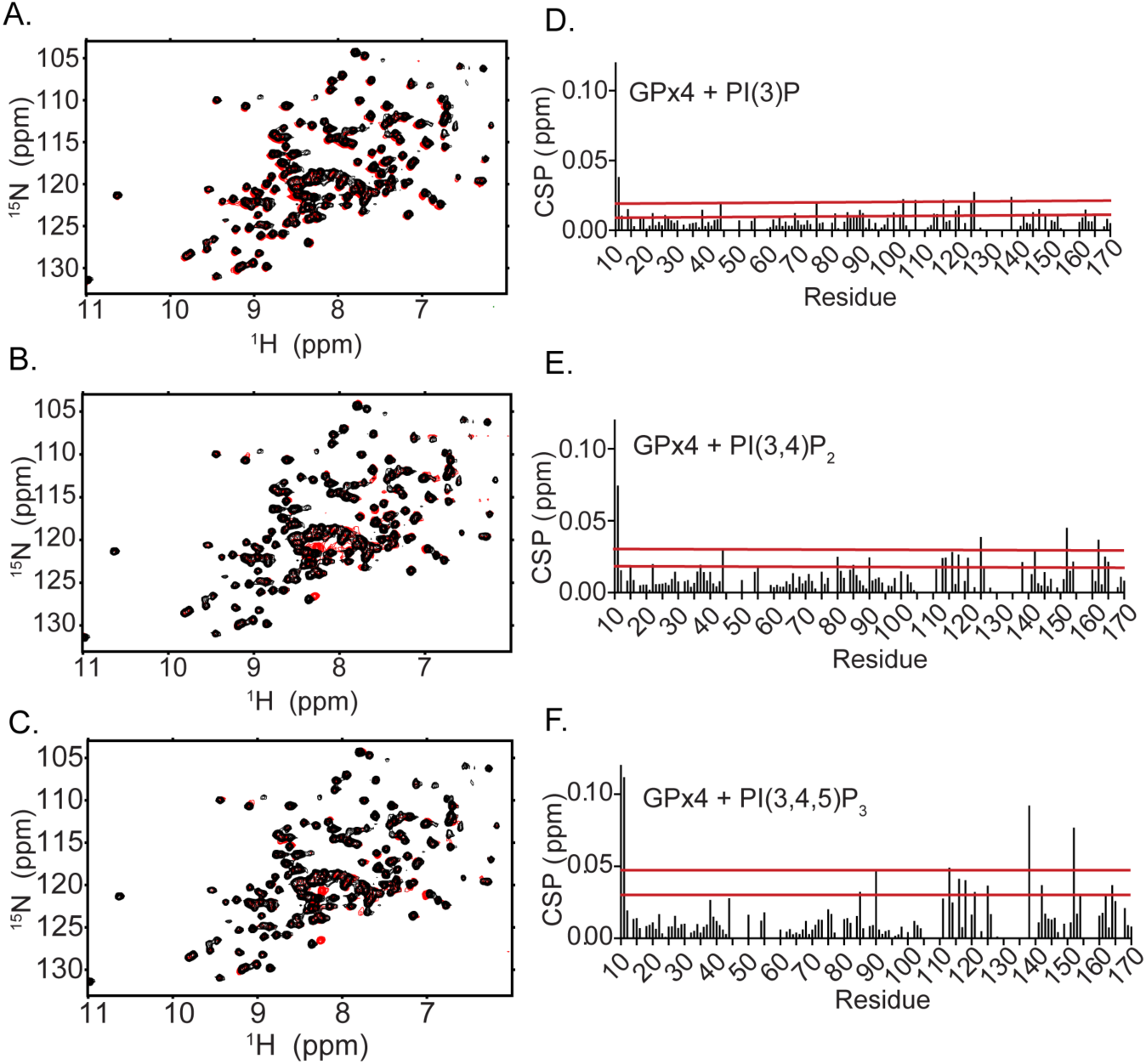
The addition of phosphorylated PIP lipids to DPC micelles causes resonance shifting in and around the cationic region of GPx4. **A.** ^1^H-^15^N HSQC of 126 µM GPx4 in 25 mM DPC micelles (black) overlaid with the spectrum of 126 µM GPx4 bound to micelles containing 25 mM DPC and 500 µM PI(3)P diC16 (red). **B.** ^1^H-^15^N HSQC of 126 µM GPx4 in 25 mM DPC micelles (black) overlaid with the spectrum of 126 µM GPx4 bound to micelles containing 25 mM DPC and 500 µM PI(3,4)P_2_ diC16 (red). **C.** ^1^H-^15^N HSQC of 126 µM GPx4 in 25 mM DPC micelles (black) overlaid with the spectrum of 126 µM GPx4 bound to micelles containing 25 mM DPC and 500 µM PI(3,4,5)P_3_ diC16 (red). **D.** Chemical shift perturbations (CSP) between GPx4 in DPC micelles and GPx4 in DPC micelles with PI(3)P diC16 with 20% trimmed mean (0.006 ppm) plus 1σ (0.006 ppm) and 2σ (0.013 ppm) (red lines). **E.** CSPs between GPx4 in DPC micelles and GPx4 in DPC micelles with PI(3,4)P_2_ diC16 with 20% trimmed mean (0.008 ppm) plus 1σ (0.012 ppm) and 2σ (0.023 ppm) (red lines). **F.** CSPs between GPx4 in DPC micelles and GPx4 in DPC micelles with PI(3,4,5)P_3_ diC16 with 20% trimmed mean (0.009 ppm) plus 1σ (0.019 ppm) and 2σ (0.038 ppm) (red lines).

Assignment of water-solubilized U46C-GPx4 resulted in 96% coverage of non-proline residues (see methods). The higher degree of coverage compared to DPC-micelle bound GPx4 served as motivation to test water-soluble analogs of PIP lipids for binding, to verify their interactions and to produce a more complete headgroup interaction surface on GPx4. IP_4_ (1,3,4,5), the headgroup analog of PI(3,4,5)P_3_, was titrated into 124 μM GPx4 and an apparant K_d_ of 980 ± 204 μM was calculated (**Figure 4A-B**). Selection of resonances for global K_d_ fitting is detailed in the materials and methods. These residues included C10-A11, H15, and H114 in the site adjacent to the membrane interaction surface and G128 and A133 within the membrane interaction surface. These interacting residues generally match well with the interaction region observed upon GPx4 binding the DPC micelles with PI(3,4,5)P_3_ diC16 incorporated. Differences between residues observed to shift with IP_4_ compared to PI(3,4,5)P_3_ diC16 in DPC micelles may be attributed to slightly altered interactions due to the presence of the micelle membrane model. Phytic acid (IP_6_ (1,2,3,4,5,6)) was titrated into GPx4 and an apparant global K_d_ was calculated (K_d_ of 314 ± 42 μM) (**Figure 4C-D)**. While not analogous to a lipid headgroup, the higher binding affinity of IP_6_ may be accounted for from the increased anionic charge from two additional phosphorylations. The residues that showed the greatest shifting upon IP_6_ titration are positioned adjacent to the membrane binding site in a similar area to the IP_4_ binding and include C10-A11, A113, L116, and K118 with I129 in the binding site having significant CSPs. IP titrations were also performed with IP_1_ (1), IP_3_ (1,3,4), IP_3_ (1,3,5) and IP_3_ (1,4,5), analogs of PI and PIP_2_ lipids, with minimal shifting observed, demonstrating that they have a significantly weaker binding affinity (**Supplementary Figs. S10-13**).

**Figure 4.**
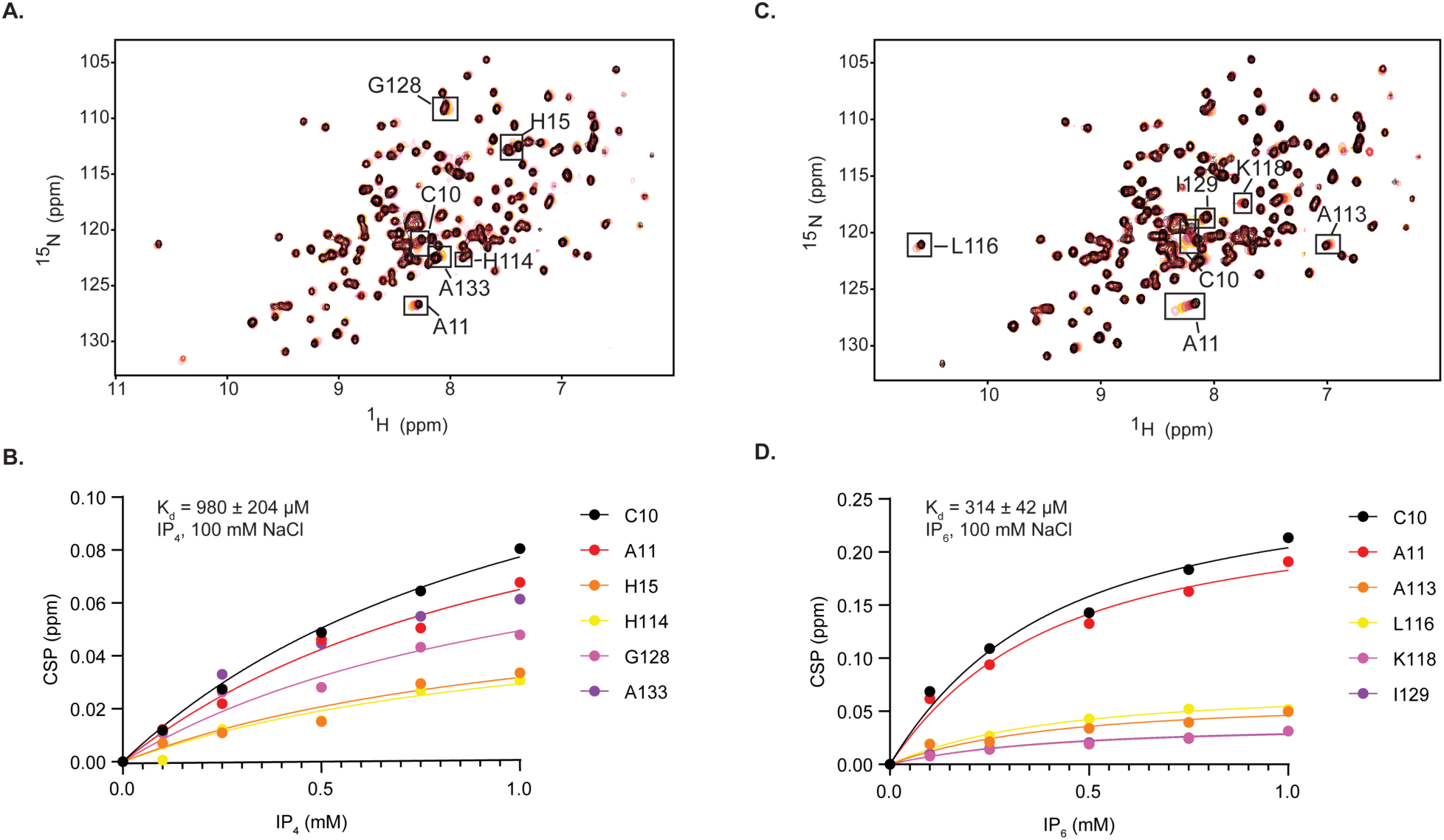
Titration of IP_4_ and IP_6_ in 124 μM GPx4 allows calculation of apparent K_d_s. **A.** ^15^N-HSQC overlay of apo GPx4 (pink) titrated with IP_4_ (1,3,4,5), within buffer containing 100 mM NaCl, in the following sequence: 1 mM IP_4_ (black), 0.75 mM IP_4_ (red), 0.5 mM IP_4_ (magenta), 0.25 mM IP_4_ (coral), 0.1 mM IP_4_ (gold). **B.** Plot of representative CSPs used to extract an apparent K_d_ value for IP_4_, K_d_ = 980 ± 204 μM. Selection of resonances for global K_d_ fitting is detailed in the materials and methods. **C.** ^15^N-HSQC overlay of apo GPx4 (pink) titrated with IP_6_, within buffer containing 100 mM NaCl, in the following sequence: 1 mM IP_6_ (black), 0.75 mM IP_6_ (red), 0.5 mM IP_6_ (magenta), 0.25 mM IP_6_ (coral), 0.1 mM IP_6_ (gold). **D.** Plot of representative CSPs used to extract an apparent K_d_ value for IP_6_, Kd = 314 ± 42 μM. The resonances selected were the top shifting residues with 1 mM IP_6_. Selection of resonances for global K_d_ fitting is detailed in the materials and methods.

To understand whether electrostatics is a major driver of PIP-headgroup interactions, as previously implicated in other lipid-GPx4 interactions,^20,21^ IP_4_ was titrated in buffer containing higher (500 mM) and lower (50 mM) NaCl concentrations. Varying the ionic strength of solutions is useful for understanding the electrostatic contributions of intermolecular interactions.^49,50^ In buffer containing 50 mM NaCl, the apparent affinity increased to a K_d_ of 309 ± 110 μM (**Figure 5A-B**), a three-fold enhancement in affinity compared to IP_4_ binding in buffer containing 100 mM NaCl. We note that this titration was only conducted up to an IP_4_ concentration of 0.25 mM, beyond which aggregation effects were observed. Conversely, when IP4 was titrated against GPx4 in buffer containing 500 mM NaCl, no significant GPx4-IP_4_ interaction was observed (**Figure 5C-D**). Considering the weakened affinity at higher ionic strength and the enhanced affinity at a lower ionic strength, these results point to electrostatics being a major driver of the interaction between GPx4 and PIP headgroups.

**Figure 5.**
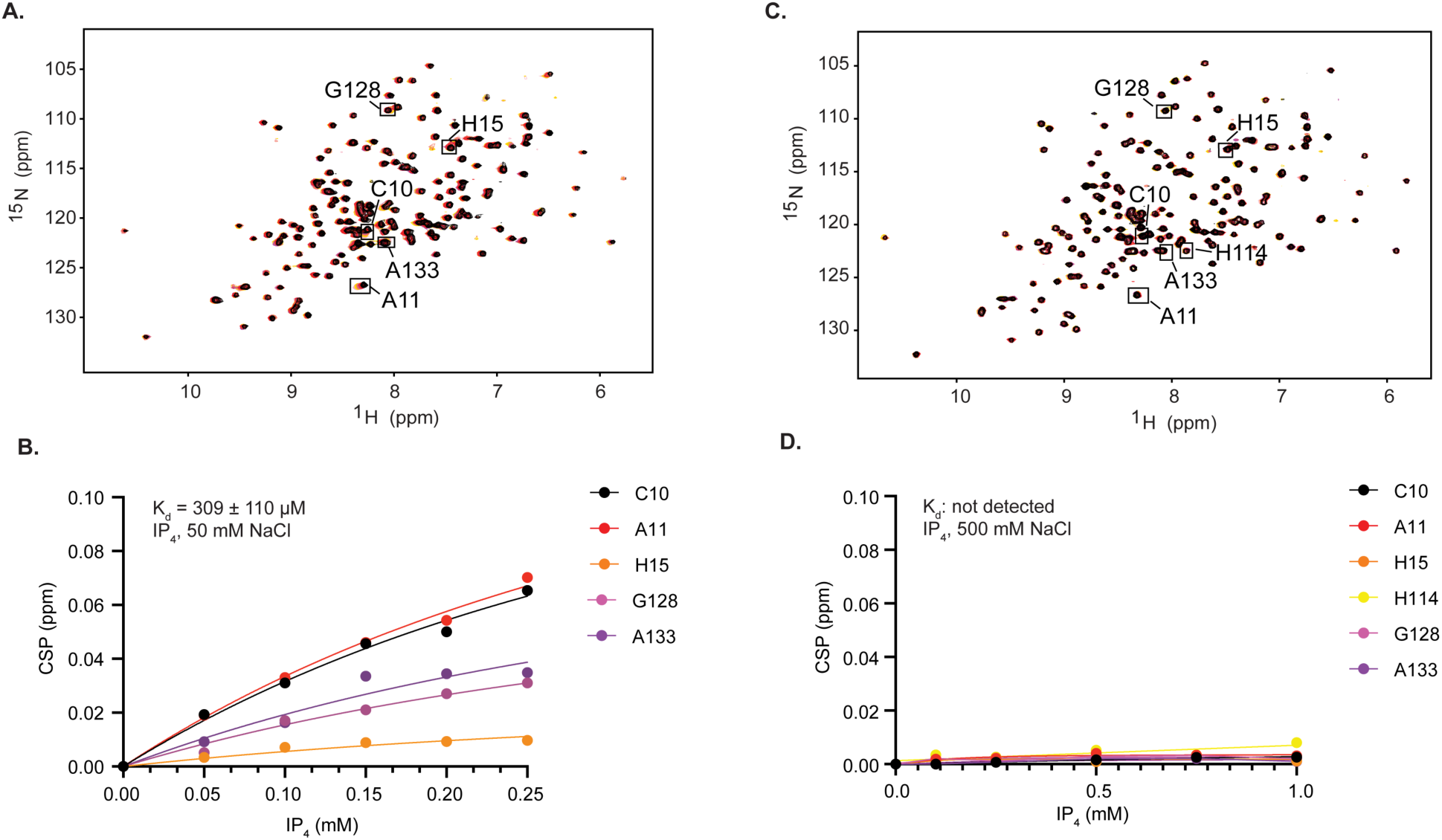
Titration of IP_4_ with varying NaCl concentration reveals an electrostatic component. **A.** ^15^N-HSQC overlay of apo GPx4 (pink) titrated with IP_4_ (1,3,4,5) in buffer containing 50 mM NaCl, in the following sequence: 0.25 mM IP_4_ (black), 0.20 mM IP4 (red), 0.0.15 mM IP_4_ (magenta), 0.10 mM IP_4_ (coral), 0.05 mM IP_4_ (gold). **B.** Plot of representative CSPs used to extract an apparent K_d_ value for IP_4_, K_d_ = 309 ± 110 μM. Selection of resonances for global K_d_ fitting is the same as used in Figure 4; note that H114 is excluded here due to resonance overlap in these buffer conditions. **C.** ^15^N-HSQC overlay of apo GPx4 (pink) titrated with IP_4_ (1,3,4,5) in buffer containing 500 mM NaCl, in the following sequence: 1 mM IP_6_ (black), 0.75 mM IP_6_ (red), 0.5 mM IP_6_ (magenta), 0.25 mM IP_6_ (coral), 0.1 mM IP_6_ (gold). **D.** Plot of representative CSPs through the titration, revealing minimal interaction of IP_4_ with GPx4 under high-NaCl conditions. Displayed residues are the same as used in Figure 4.

To better understand the structural determinates of PIP lipid interactions, crystallographic data was collected for the IP_4_ (1,3,4,5)-GPx4 complex (**Figure 6, Supplemental Table S1,** PDB ID: 35WP). GPx4 crystallized as a tetrameric structure with only monomers C and D binding to IP_4_. IP_4_ sits in a shallow pocket in between two loops, the previously described ‘fin-like’ loop,^19^ which plays a significant role in membrane engagement, and the opposite loop containing N109-D111. Monomers A and B do not have IP_4_ binding due to the site being occupied by D111 of the symmetrical units. Using the PPM 3.0 server^51^ and MaSIF-PMP^52^, the membrane-interface prediction confirms the feasibility of the location and orientation of the lipid headgroup against the membrane model with the overlay of the crystal structure matching the PPM 3.0 server results (**Figure 6A, Supplementary Fig. S14**). Here, the IP_4_ is positioned as if a PI(3,4,5)P_3_ headgroup protrudes from the membrane surface, which is expected for PIP lipids.^53^ The IP_4_ is positioned so that the 1-phosphate and the 5-phosphate interact closely with the hydrophobic fin-loop in the membrane interacting site, specifically N132 and A133. 1-phosphate, analogous to the phosphodiester linker in PIP_3_, and is positioned so that the lipid tail, in a biological context, would be oriented towards the active site of GPx4, suggesting the headgroup binding positions the hydroperoxidated tails for catalytic processing (**Figure 6B**). Examination of the interactions between ligand and protein in chain C reveal specific hydrogen bonding to IP_4_ comes from the side-chain oxygen of N109, backbone nitrogen and oxygens of G110, side-chain oxygen and backbone nitrogen of N132, and the backbone nitrogen of A133. (**Figure 6C-D**). The residues interacting with IP_4_ can be mapped as a surface proximal to the cationic interacting site of GPx4. This binding region confirms the site anticipated from NMR results with the same amino acids participating in the IP_4_ binding. Large CSPs are observed by NMR in the N-terminal tail upon IP_4_ titration, but crystallographic structure does not show direct interactions with the ligand nor conformational changes. This may be due to a different IP_4_-induced conformation in solution that is not captured via crystallography.

**Figure 6.**
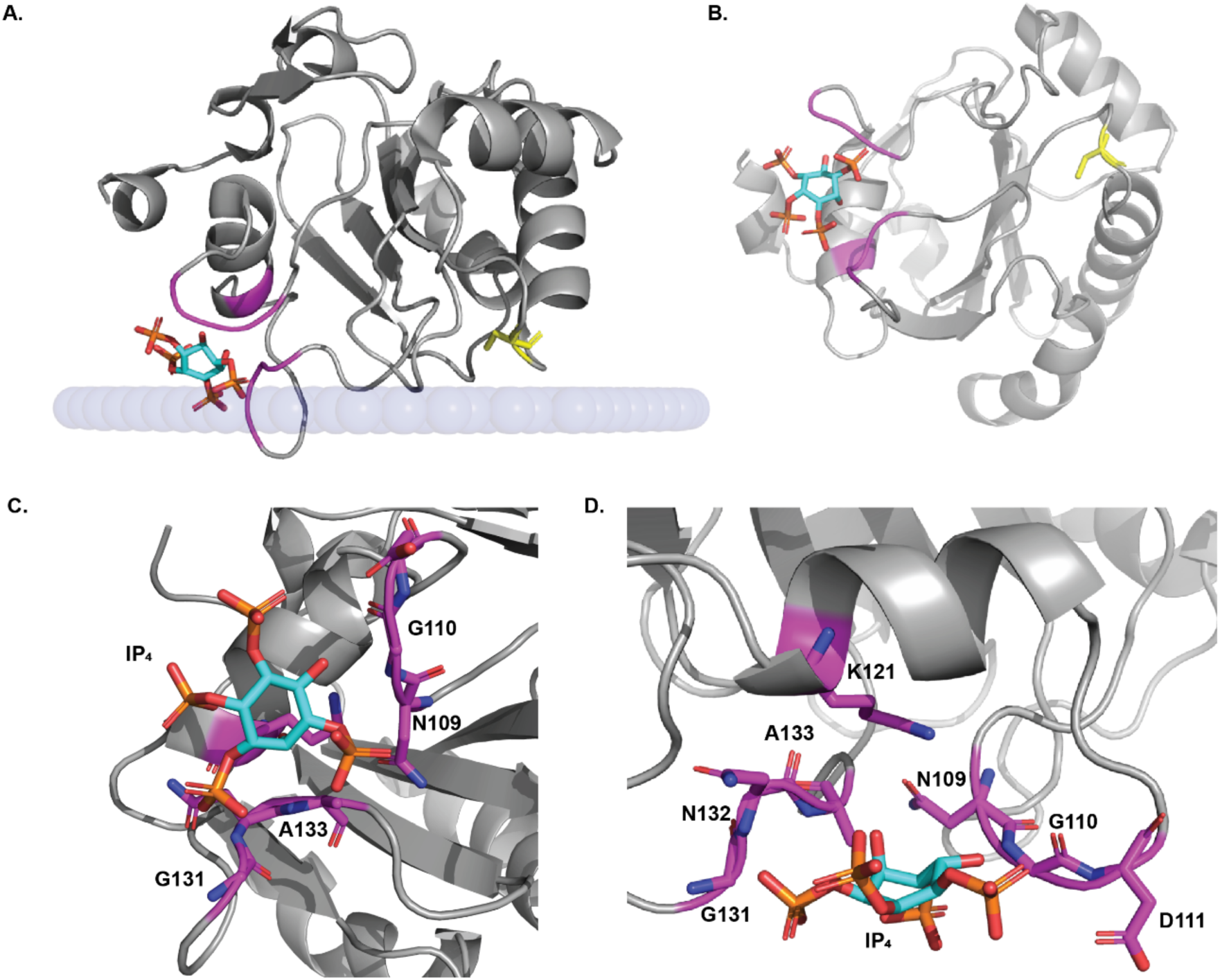
GPx4 co-crystalized with IP_4_. **A.** Crystal structure of GPx4 (gray cartoon) with IP_4_ bound against a simulated membrane layer (blue spheres) determined through PPM 3.0 server using PDB ID 2OBI. Magenta highlights residues interacting with IP_4_ include N109-D111, K121, and G131-A133 and C46 is displayed as yellow sticks. **B.** 90° rotation of the complex structure, with a view from membrane bilayer. **C-D.** Zoomed in orientations of IP_4_ bound to GPx4 with interacting residues (labeled in figure) in magenta sticks with nitrogen atoms in blue and oxygen atoms in orange. IP_4_ ring is in cyan with oxygen atoms in orange and phosphorus atoms in yellow. Hydrogen atoms not shown.

We performed molecular dynamics (MD) simulations to relate the observed IP_4_ binding site, referred to hereafter as ‘hotspot residues’ (**Supplementary Table S2**), to potential interactions between GPx4 and PI(3,4,5)P_3_ and how these interactions may relate to GPx4’s function. To evaluate whether the IP_4_ conformation with GPx4 is compatible with the catalytic reduction of a lipid tail, we performed MD simulations of a GPx4–PI(3,4,5)P_3_ complex in bulk water. The starting configuration for these simulations had the head group of a full PIP molecule aligned to the IP_4_-bound co-crystal structure (**Figure 7A, Supplementary Fig. S15**). The GPx4-PI(3,4,5)P_3_ complex was stable throughout the simulations with no unbinding observed. We tracked the minimum distance between the center-of-mass (COM) of the 40s loop, which is known to play critical roles in GPx4’s membrane binding and catalytic activity^20^, and the terminal groups of the PI(3,4,5)P_3_ lipid tails (**Figure 7B**). This demonstrates that the 40s loop makes frequent contact with the terminal groups of both PI(3,4,5)P_3_ lipid tails, confirming the observed IP_4_-GPx4 co-crystal structure provides a robust structural scaffold supporting the spatial orientation required for tail hydroperoxidase activity.

**Figure 7.**
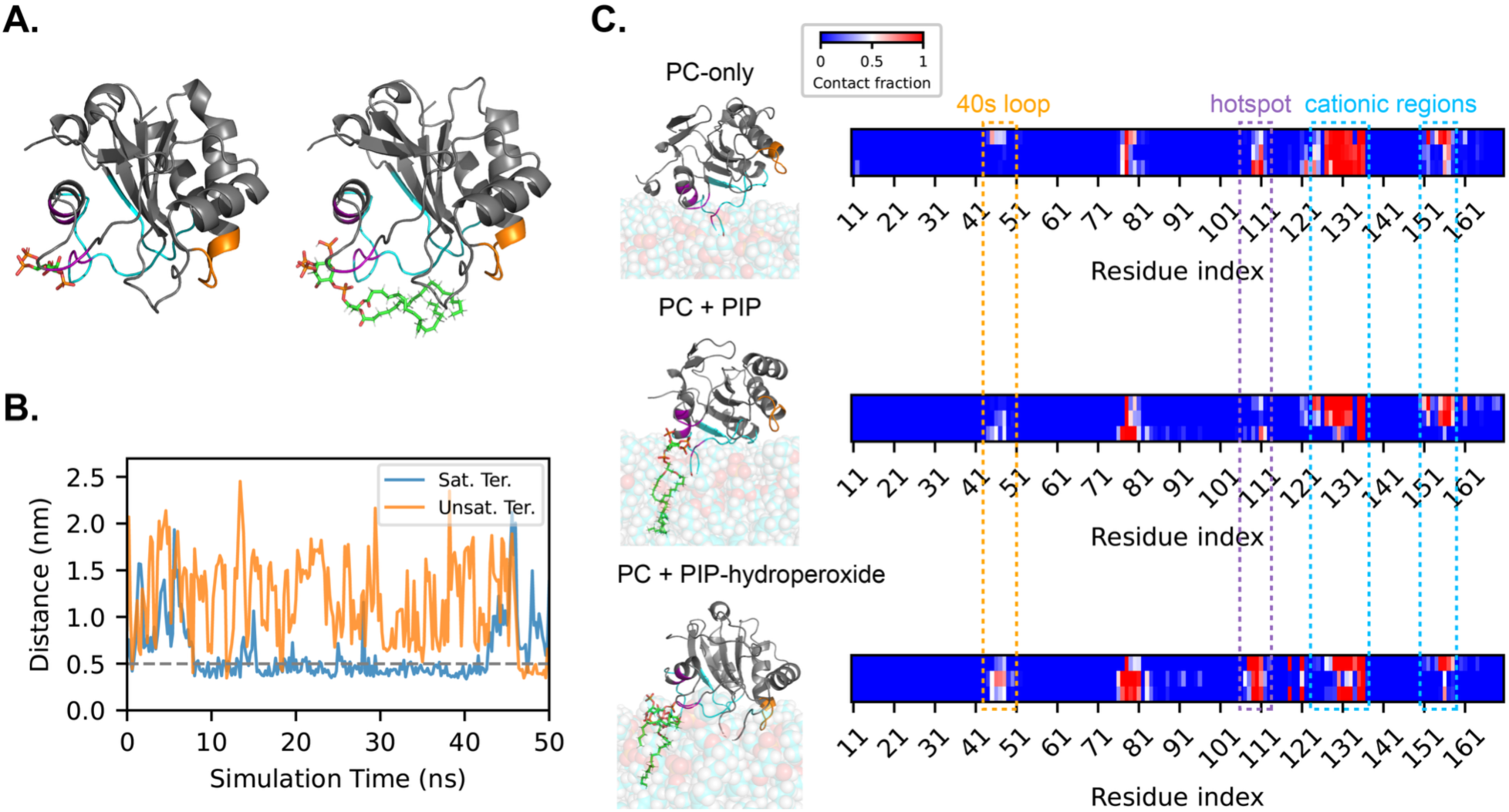
MD simulations sampling GPx4 interactions with PI(3,4,5)P_3_. **A.** Co-crystallized structure of IP_4_-bound GPx4 (left) and a representative snapshot of the PI(3,4,5)P_3_-bound GPx4 structure prepared for MD simulations (right). GPx4 is shown in a gray ribbon representation, with the 40s catalytic loop highlighted in orange, cationic regions in cyan, L130 in red, and hotspot residues in magenta. IP_4_ and PI(3,4,5)P_3_ are shown in stick representations with carbon atoms in green, hydrogen atoms in white, oxygen atoms in red, and phosphorus atoms in orange. Solvent molecules are not shown. **B.** Distances between the two PIP lipid termini and the center-of-mass of the GPx4 40s loop during a 50 ns unbiased simulation of PI(3,4,5)P_3_-bound GPx4. The trajectory from one simulation replicate is shown; trajectories for two additional replicas are in **Supplementary Figure S16**. “Sat. Ter.” denotes the distance between the GPx4 40s loop and the saturated lipid terminus of PIP and “Unsat. Ter.” denotes the corresponding distance for the unsaturated lipid terminus. Distances within 0.5 nm are considered contacts, as indicated by the gray dashed line. **C.** Barcode plots showing GPx4–lipid contacts across different membrane systems: DOPC-only, DOPC + PI(3,4,5)P_3_, and DOPC + PI(3,4,5)P_3_-hydroperoxide. Representative snapshots of membrane-bound GPx4 are shown for each system. GPx4 and PIP lipids are represented as in panel **A.** DOPC lipids are shown in a transparent van der Waals representation using the same color scheme as PI(3,4,5)P_3_ except that carbon atoms are cyan. Solvent molecules are not shown to aid visualization. For the barcode plots, time-averaged contact fractions for GPx4 residues are shown using a blue-to-red color scale ranging from 0 to 1, with a value of 1 indicating the residue is in contact with the membrane for the entire trajectory. Each row corresponds to one of three replicas for each membrane system. The orange dashed box marks residues within the 40s loop, the magenta box marks residues adjacent to hotspot residues, and cyan boxes mark residues within cationic regions. U46 and W136 of the catalytic triad are located in the 40s loop and cationic regions, respectively, while Q81 is labeled in the plot.

We examined the effect of PI(3,4,5)P_3_ on GPx4 membrane binding, and the specific residues in contact with the membrane, by simulating GPx4 in three different membrane systems: pure PC, PC with a single PI(3,4,5)P_3_ lipid, and PC with a single PI(3,4,5)P_3_ lipid with a hydroperoxidized tail (**Figure 7C; Supplementary Figs. S17-19**). DOPC was selected as a model PC lipid for these simulations because it forms a fluid lipid bilayer with interfacial chemical properties comparable to those of DPC (used in the NMR experiments) and it has previously been employed in experimental studies characterizing GPx4–PC membrane interactions.^20^ Initial configurations for the simulations were prepared by applying biasing forces to move the protein into contact with the membrane (**Supplementary Fig. S18**); in the case of membranes with PI(3,4,5)P_3_ (with and without a hydroperoxidized tail), the protein was moved such that the PI(3,4,5)P_3_ head group was aligned with the IP_4_ binding site from the co-crystal structure. The production runs without the biasing forces showed GPx4 binding to the membrane and interactions with the PI(3,4,5)P_3_ head group to be stable throughout the simulation, permitting analysis of GPx4 residues in contact with the membrane. **Figure 7C** shows time-averaged residue-membrane contacts, with a contact counted when the distance between a residue and lipid heavy atom was less than 0.5 nm. In addition to monitoring functional regions such as the 40s loop and hotspot residues, we tracked the dynamics of the cationic regions.^20^ In pure PC bilayers, the protein exhibited stable interactions mediated primarily by contact between the cationic regions and membrane phosphate headgroups (**Supplementary Fig. S19**). However, the introduction of PI(3,4,5)P_3_ induced a distinct shift in binding dynamics: while total interactions via cationic regions decreased, membrane contacts involving the 40s loop and residues proximal to the catalytic triad member Q81 significantly increased. This trend was amplified in the presence of peroxidized PI(3,4,5)P_3_, suggesting that GPx4 binding to the PI(3,4,5)P_3_ hydroperoxide actively promotes catalytic contact. To further investigate contributions of GPx4 cationic residues to stable PI(3,4,5)P_3_ binding and overall membrane association, we quantified the propensity of cationic residues to contact the PI(3,4,5)P_3_ headgroup and/or the membrane surface excluding PI(3,4,5)P_3_ using the same contact criterion and trajectories as in **Figure 7C**. Key cationic residues contributing to stable PI(3,4,5)P_3_ and membrane association were identified as those maintaining contact for more than 50% of the analyzed simulation time. **Supplementary Figure S20** shows that cationic residues located within the membrane binding regions (i.e., 40s loop, Q81-adjacent region, cationic region) predominantly interacted with PI(3,4,5)P_3_ and the membrane surface, thereby contributing to stable GPx4–membrane association. Consistent with the interaction patterns discussed above, the introduction of PI(3,4,5)P_3_ into the PC membrane increased contacts involving the 40s loop and residues proximal to the catalytic triad member Q81 (**Supplementary Figs. S20-21**). Ultimately, these trajectories indicate that the PIP headgroup serves as a critical conformational scaffold that stabilizes the complex and optimally orients the catalytic regions toward the membrane interface.

Finally, to quantify the accessibility of the lipid tail to catalysis, we employed umbrella sampling (US) to compute the potential of mean force (PMF) for the protrusion of unsaturated PI(3,4,5)P_3_ lipid tails from the DOPC bilayer toward the GPx4 40s loop (**Figure 8**). The PMF quantifies the free energy change associated with a varying reaction coordinate. Simulations were initiated from the same configurations used for the results in **Figure 7C** but with biasing forces to position the terminus of the unsaturated PI(3,4,5)P_3_ lipid tail at different distances from the center-of-mass of the 40s loop (*d_COM_*). Using *d_COM_* as a reaction coordinate, we computed the free energy associated with moving the tail into contact with the 40s loop (corresponding to *d_COM_* = 0.5 nm) from its initial equilibrium position in the membrane, defined as Δ*G_protusion_*. For the GPx4-bound conformation, the free energy cost for the protrusion of a peroxidized lipid tail was substantially lower (16.59 ± 0.59 kcal/mol) than that of its native counterpart (31.03 ± 1.03 kcal/mol). No change in the membrane-bound GPx4 conformation was observed, but the equilibrium state of the peroxidized PI(3,4,5)P_3_ tail was found to correspond to a value closer to the 40s loop, which we attribute to the increased tail hydrophilicity upon hydroperoxidation which positions the tail closer to the GPx4-membrane interface.

**Figure 8.**
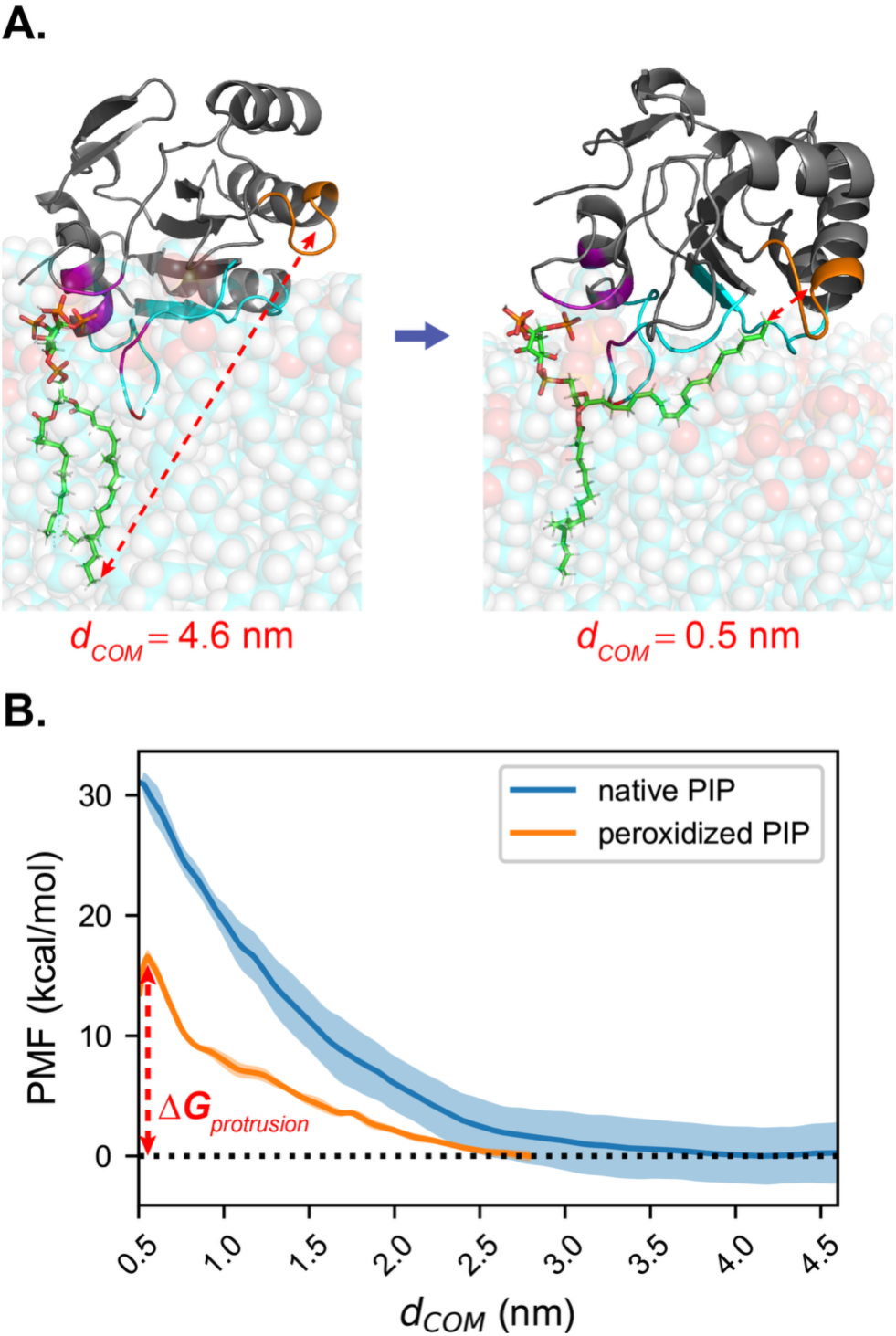
Comparison of protrusion behavior of the unsaturated PI(3,4,5)P_3_ lipid terminus as a function of peroxidation state. **A.** Representative snapshots illustrating the protrusion of the native PI(3,4,5)P_3_ unsaturated lipid tail toward the GPx4 40s loop. The process begins from a stably membrane-inserted PIP configuration, followed by translocation of its unsaturated lipid terminus across the DOPC bilayer toward the GPx4 40s loop. The color scheme follows the same definition used in Figure 7C. Red dashed arrows and corresponding annotations indicate the distance between the center-of-mass of the GPx4 40s loop and the unsaturated lipid terminus of PI(3,4,5)P_3_, *d_COM_*. **B.** Potential of mean force (PMF) as a function of *d_COM_*. The state in which the unsaturated PIP lipid terminus is in contact with the GPx4 40s loop corresponds to *d_COM_* = 0.5 nm, whereas the most equilibrated membrane-inserted states correspond to *d_COM_* = 4.6 nm and 2.8 nm for native and peroxidized PI(3,4,5)P_3_, respectively. Solid lines and shaded regions represent the average and standard deviation of converged PMFs across replicas for each system. The protrusion free energy, Δ*G_protrusion_*, was 31.03 ± 1.03 kcal/mol for native PI(3,4,5)P_3_ and 16.59 ± 0.59 kcal/mol for peroxidized PI(3,4,5)P_3_.

## Discussion

The work presented here suggests that PIP lipids, particularly the tris-phosphorylated species, act as preferred substrates for GPx4, alongside known anionic lipid binders such as CL, PS, PA, and others.^21^ This high affinity was expected due to the presence of a cationic site located within the membrane binding and lipid interaction surface of GPx4, making negatively charged lipids preferential binders. Previously, electrostatically driven affinity of GPx4 for negatively charged lipids and membranes has been demonstrated, leading to the conclusion that the lipid type and overall cationic character of the binding site both contribute to the GPx4-membrane interaction. Characterizing interactions with highly anionic PIP lipids allows further understanding of GPx4 binding and structure in a biological context since PIP lipids are enriched in the inner leaflet of the plasma membrane, where the cytosolic isoform of GPx4 has ample access.^27^ NMR analysis showed an increase in resonance shifting as the number of phosphorylations increased, importantly, in and near the cationic and membrane-binding site. This confirms that higher anionic charge results in a greater interaction between GPx4 and PIPs and begins to map a headgroup-specific interacting site. By comparing the binding sites of the phosphorylated PIPs with and without a membrane model with the CSP mapping of the IP_4_ and the crystal structure, there appears to be increased shifting in a region directly adjacent to the cationic membrane interacting site. This indicates a binding site for lipid headgroups that is partially overlapped with the known cationic membrane interacting site based, including residues N109-D111 and G131-A133. Characterizing the interaction between IP_4_ and GPx4 in buffers in varying ionic strengths indicates that electrostatics is likely a major contributor to GPx4 binding to PIPs.

This specific binding site modifies the current conformation of GPx4-lipid interactions from a broad cationic surface while retaining the ability to reduce lipid hydroperoxides and allows us to rationalize the apparent higher affinity GPx4 has for PIPs. The headgroups of PIP lipids protrude out from the membrane, resulting in binding along the side of the protein, in the region adjacent/partially within the cationic membrane interacting site. The lipid tail is predicted to be long enough (∼18 Å) to wrap around the protein within the membrane and allow for the unsaturated tail to reach the catalytic triad, a behavior confirmed by MD simulations of GPx4-PIP interactions in both solution and in a PC membrane. This relationship not only explains a preference for PIP lipids, which are prevalent within the native membranes that GPx4 interacts with,^54^ but also further explains the anionic charges that GPx4 has a tendency to bind with and the structural underpinnings of the catalytic mechanism of GPx4. This orientation solidifies the idea that the apolar residues within the GPx4 cationic membrane interacting site are involved in the positioning of the unsaturated tail towards/within the active site suggesting an enhancement in catalytic activity.^23^ The increased contact frequency may be attributed to the pronounced tilting propensity of peroxidized PIP within the bilayer. When GPx4 is anchored to the PIP headgroup, the lipid’s tilted orientation drives a global reorientation of the protein—pivoting the 40s loop and catalytic residues toward the membrane surface in a lever-like mechanism. Free energy calculations support that this tilting reduces the thermodynamic cost for bringing hydroperoxidized PIP tails into contact with the catalytic region of membrane-bound GPx4. We attribute the reduced barrier for protrusion to the increased hydrophilicity of the unsaturated lipid tail following its peroxidation. This reduction in free energy implies that the peroxidation of a PIP lipid tail not only enhances GPx4 recruitment but may also facilitate its own reduction. While the overall free energy for protrusion remains larger than thermal energy, the reduction in this barrier facilitates a closer positioning of the lipid to the GPx4–membrane interface. This suggests that GPx4 may also act on tails from nearby PIP lipids other than the one to which it is bound, particularly if PIP lipids cluster in the membrane as suggested previously.^55–59^

## Conclusions

We have determined that PIP lipids, especially PI(3,4,5)P_3_, may serve as preferred substrates of GPx4 and provided new structural information about how GPx4 binds with lipid headgroups to perform its function. Understanding interactions and functional consequences of various lipids with GPx4 provides deeper insight into its functionality and mechanism. Ability of GPx4 to process PIP lipids may translate into membrane homeostasis and cell health. PIP lipids are abundant in the plasma membrane and are targets of hydroperoxidation by LOX enzymes, which would drive ferroptosis and disrupt PIP membrane signaling pathways if left unchecked.^54^ Compared to healthy cells, PI(3,4,5)P_3_ is enriched in a wide range of cancers due to upregulation of PI3K, which may suggest an particularly important role for GPx4 in reducing these lipids in ferroptosis-sensitive cancer cells.^38,39^ Despite the potential importance of PIP lipids, the low abundance of each individual PIP species and short lifetimes of hydroperoxidations have limited exploration of the role of PIP-hydroperoxidation in ferroptosis.^60^ These results highlight that there may be an important role for GPx4 in elimination of PIP oxidative damage and this interaction may be important for ferroptosis regulation in cells with higher PI3K activity which includes a wide range of cancer cells. In addition, enrichment of PIP lipids within the plasma and other membranes may in part drive localization of GPx4 and direct its ferroptosis protective activity. PIP lipids are vital to a variety of biological functions and understanding how GPx4 can interact with PIP lipids allows for a deeper appreciation for its ability to protect against an array of lipid hydroperoxidations and potentially preserve PIP signaling networks.

## Materials and Methods

### Protein Growth and Preparation

The synthetic gene for residues 7-170 of the human cytosolic isoform of U46G-GPx4 with a TEV cleavable poly-histidine tag in the vector pNIC28-Bsa4 was a gift from Nicola Burgess-Brown (Addgene plasmid #38797). For this work, the glycine in the 46^th^ position was mutated to a cysteine and is referred to as GPx4.^61^ Further details are presented in the **Supplementary Information**.

GPx4 was grown and prepared as previously reported.^20^ GPx4 encoding plasmid was transformed into BL21 *E. coli* cells where they were grown and harvested for lysis. The lysis and purification steps were completed through sonication and Ni-NTA affinity chromatography. Unless specified, the His-tag was then cleaved with a TEV protease and the protein was then repurified. Protein concentration and purity were assessed via Bradford assay and SDS-PAGE gel, respectfully. Complete details are available in the **Supplementary Information**.

### Lipid Overlay Assay

We adapted a microplate reader detected, time-resolved fluorescence method using europium labeled secondary antibodies, originally developed for western-blotting, for the LOAs. The LOA protocol was based on the Echelon Biosciences PIP Strips recommended protocol with some modifications made to utilize the Molecular Devices Scan Later assay kit for visualization. Breifly, the His-tagged GPx4 was incubated with the membrane strip before the 1° antibody was added followed by the Eu-labelled 2° antibody with wash steps performed between each addition. The membrane strip was then scanned on a SpectraMax iD5 Plate Reader (Molecular Devices) using Time-Resolved Fluorescence imaging with recommended settings for a Western Blot Assay. The excitation wavelength was 340 nm and the emission wavelength was 616 nm. For complete details, please see the **Supplementary Information**.

### NMR Spectroscopy and analysis

All NMR samples were prepared with ^15^N-isotopically labeled protein. The GPx4 buffer typically consisted of 20 mM Bis-Tris pH 6.0, 0.1 M NaCl, and 20 mM DTT and 10% D_2_O was used as lock solvent, with the NaCl concentrations adjusted to 50 mM or 500 mM to probe electrostatic dependence of IP_4_ binding. Please see **Supplemental Information** for micelle preparation details.

^1^H-^15^N HSQC experiments were collected on 600 MHz Bruker AVANCE III with a room temperature probe. Resonance assignment data of aqueous GPx4 was collected on 700 MHz Bruker AVANCE III with room temperature probe using the following 3D experiments: HNCA, HNCB, HN(CO)CA, and CBCACONH, resulting in 96% of non-proline resonances being assigned.^46,62^ For assignments of micell-bound GPx4, HNCA and HN(CO)CA were collected with non-uniform sampling (NUS), 25% sampling with the Bruker NUS addendum used for reconstruction. 75% of non-proline resonances were assigned for DPC-micelle bound GPx4. All experiments were collected at 25°C. NMR data was processed using NMRPipe and analyzed using NMRFAM-Sparky.^63,64^ Chemical shift perturbation calculations were completed with the following formula:

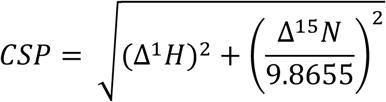

Where Δ^1^H and Δ^15^N represent the changes in the ^1^H and ^15^N chemical shifts for each resonance.

For NMR titrations, CSPs per concentration were calculated and apparent global K_d_ values were determined. To fit, individual resonances with CSPs > 0.025 at the highest titration point and an R^2^ > 0.95 from individual apparent K_d_ values were selected. Selected residues were included in a global fit and residues with R^2^ < 0.95 in the global fit were removed, with only a single residue removed here from IP_4_ and IP_6_ data sets. The final selected resonances were then all used to calculate a global fit for the apparent K_d_ value. The K_d_ calculation was made from the following equation, fitting all selected residues to a global K_d_ value:

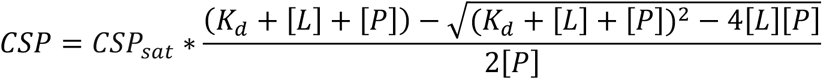

Calculations were performed in GraphPad Prism 10.0.0.

### Crystallography

Protein samples were prepared as described above. Ligand was added to 7.7 mg/mL sample at a concentration of 2 mM. Full methods, additional information, and refinement table are provided in **Supplementary Information.**

### Molecular Dynamics Simulations

3D structures of GPx4 were modeled using the structure with PDB ID 2OBI as template or the complete structure of IP_4_-bound GPx4 characterized by x-ray crystallography, the former for phosphocholine (PC)-only membrane system and the latter for GPx4-IP_4_ and PIP-introduced PC membrane systems. All proteins were prepared with standard charged termini (–NH_3_^+^, –COO^−^) and all residues were protonated at pH 6.0 to match the conditions used in the experiments. All PC membrane systems were modeled using 1,2-dioleoyl-sn-glycero-3-phosphocholine (DOPC) lipids using the *Membrane Builder* module of CHARMM-GUI.^65,66^ The PIP lipid was modeled using the PI(3,4,5)P_3_ structure from the CHARMM Small Molecule Library of CHARMM-GUI Archive^66^ and the PIP hydroperoxide structure was modeled using Avogadro v1.2.0^67^ and CHARMM-GUI *Ligand Reader & Modeler* module (**Supplementary Fig. S17**).^66,68^ All systems were neutralized with 0.10 M NaCl to match the experimental conditions of the previous NMR study. All systems were modeled using the CHARMM36m forcefield with WYF parameter for cation-π interactions^69–71^ and the TIP3P water model. Molecular structures and force field parameters for GPx4, PIP lipids, and the lipid bilayer were generated using the CHARMM-GUI Input Generator.^65,66,68,72,73^ The forcefield parameters for PIP hydroperoxides were refined following the approach of Hu et al. 2025^74^ to ensure charge consistency with the native lipid species. Complete system preparation details are presented in the **Supplementary Information**.

All simulated systems were first energy minimized using steepest descent algorithm until the maximum force between atoms reached the criterion of < 1000 kJ mol^−1^ nm^−2^. All equilibration and production simulations of systems were performed under NPT conditions at 298.15 K using a velocity-rescale thermostat (time constant = 1.0 ps) and at 1 bar using a stochastic cell-rescale barostat (time constant = 5.0 ps, compressibility = 4.5×10^−5^ bar^−1^). All MD simulations were conducted with a 2 fs timestep using the leapfrog integrator in GROMACS 2021.^75^ Verlet lists were generated with a 1.2 nm neighbor list cutoff, van der Waals interactions were modeled using a Lennard-Jones potential with a 1.2 nm cutoff that was smoothly shifted to zero between 1.0 and 1.2 nm, and electrostatic interactions were calculated using the smooth particle-mesh Ewald method with a short-range cutoff of 1.2 nm.^76^ Bonds involving hydrogen atoms were constrained using the LINCS algorithm.^77^ All the simulations presented in this study were performed in three replicas starting from different initial velocities. Complete simulation details are provided in the **Supplementary Information**.

## Supplementary Material Description

The **Supplementary Information** provides spectral overlay for mentioned ligands, as discussed in **Results**. Additionally, full methods and analysis are provided for crystallography and molecular simulations, with the corresponding supplemental figures and tables.

## Acknowledgements

This research was supported by the National Institute of General Medical Sciences of the National Institutes of Health under Award Number R35GM147221 to B.F. This content is solely the responsibility of the authors and does not represent the official views of the National Institutes of Health. We gratefully acknowledge support from the Eli Lilly and Company 2024 ACACC Young Investigator award to B.F. This material is based upon work supported by the National Science Foundation under Grant No. 2245375 to R.C.V.L. We gratefully acknowledge technical assistance from Dr. Yun Qu.

## Author Contributions

SHW: Methodology; Visualization; Writing – original draft; Writing – review & editing; Investigation. BP: Methodology; Visualization; Writing – original draft; Writing – review & editing; Investigation. RSL: Investigation; Visualization; Writing – review & editing. CLL: Methodology; Writing – original draft; Writing – review & editing; Investigation. FM: Investigation; Validation; Data Curation; Writing – original draft; Writing – review & editing; RCVL: Conceptualization; Funding acquisition; Project administration; Supervision; Writing – original draft; Writing – review & editing. BF: Conceptualization; Funding acquisition; Project administration; Supervision; Writing – review & editing.

## Additional Information (incl. Competing Interests Statement)

The authors declare no competing interests.

