## Supplemental Information for "Revealing interactions between glutathione peroxidase 4 and phosphoinositides"

**Supplementary Methods**

**Protein Growth and Preparation**

The Agilent QuikChange II XL Site-Directed Mutagenesis Kit was used per manufacturer's instructions, and the resulting DNA was transformed into *XL10-Gold E.coli*. The following day, a colony from the transformation plate was added to 10 mL of LB media in the presence of kanamycin and grown overnight. A Qiagen QIAprep Spin Miniprep Kit was used to purify the DNA and then sequenced by Eurofins Genomics to confirm mutation prior to use in experiments.

Briefly, the GPx4 encoding plasmid was transformed into BL21 *E.coli* cells and an overnight starter culture was used to seed 1 L cultures in shake-flasks. Protein expression was induced with isopropyl-β-D-1-thiogalactopyranoside (IPTG) at an $OD_{600}$ ~ 0.8. Cells were harvested through centrifugation, and the protein was resuspended in lysis buffer [0.1 M Tris pH 7.4, 0.3 M NaCl, 0.5 % v/v Triton, 0.1 mg/mL lysozyme, protease inhibitor cocktail, and 5 mM dithiothreitol (DTT)]. The cells were then lysed with probe sonication. GPx4 was purified via Ni-NTA affinity chromatography and then dialyzed overnight in NMR buffer [20 mM Bis-Tris pH 6.0, 0.1 M NaCl, and 20 mM DTT]. His-tagged GPx4 used for the lipid overlay assays was not processed further. For other experiments, the His-tag was cleaved by adding TEV protease to the elution fraction and the protein was dialyzed. After His-tag proteolysis, the protein was repurified using Ni-NTA resin to remove TEV, uncleaved protein, and cleaved His-tags, then dialyzed overnight into NMR buffer. Protein was quantified by Bradford assay and purity was assessed by SDS-PAGE.

**Lipid Overlay Assay**

All assays were performed using the same method at room temperature and incubated with gentle shaking. First, the strip was blocked via incubation for 1 h with 10 mL of the Molecular Devices 1x blocking buffer. Blocking buffer was removed and an additional 10 mL of 1x blocking buffer was added along with 1 μg/mL of protein (His-tagged GPx4) and incubated for 1 h. The excess buffer was removed and three washes of 5 mL with 1x Molecular Devices washing buffer were performed for 10 minutes each. Next, the primary antibody (2 mg/mL 6x-His Tag Polyclonal Antibody from Invitrogen) was added at a 1:1000 molar ratio with 5 mL 1x blocking buffer and incubated for 1 h. Three additional washes were performed. Finally, the secondary antibody (0.1 mg/mL Eu-Labeled Goat Anti-rabbit antibody from Molecular Devices) was added at a 1:500 molar ratio with 5 mL 1x blocking buffer and allowed to incubate for 1 h. An additional three washes were performed then the membrane was allowed to dry before scanning on a SpectraMax iD5 Plate Reader (Molecular Devices) using Time-Resolved Fluorescence imaging with recommended settings for a Western Blot Assay. The excitation wavelength was 340 nm and the emission wavelength was 616 nm.

**Micelle Preparation**

For the formation of the DPC micelles, 25 mM of DPC was weighed out and added to a 500 μL protein sample in NMR buffer. It was vortexed until visual clarity was reached and there were no solids present. For micelles which included PIPs, solvent solubilized lipid in 1:1 chloroform: methanol (v/v) was measured into a vial and solvent removed under nitrogen and then overnight in the speed vacuum. The DPC was then weighed and added into the vial in its powder form with the PIP lipid and formation of micelles continued as described above. Successful sample construction was confirmed via NMR.

### Crystallography

Crystallization of the GPx4-IP4 complex (PDB ID: 35WP) was carried out by the sitting drop vapor-diffusion method using a Crystal Gryphon Robot (Art Robbins Instrument) by screening against several crystallization screening kits: PEGRxHT, NatrixHT (Hampton Research), Wizard Classic 3&4 (Rigaku), and MCSG4 (Anatrace). The best crystals obtained were from reservoir solutions containing 100mM HEPES, pH 7.5, 20% PEG4000 and 10% Isopropanol. To improve the quality and size of crystals hanging drop vapor diffusion method was performed up to microliter range in 24 well VDX crystallization plates (Hampton Research). The crystal belonged to the space-group $P2_1$ with unit cell dimensions a = 76.91Å, b = 61.88 Å, c = 80.80 Å, β = 113.40 (°). Crystals were protected in reservoir solution containing 30% Isopropanol and flash-frozen in the liquid nitrogen stream. X-ray diffraction data was collected at 100K using a Rigaku Micro-Max-007 HF X-ray generator and EIGER 4M detector. The data was processed with CrysAlis*Pro* 41.64.122a (Rigaku) and CCP4 suite of programs.[1]

The structure was solved with the *Phaser-MR* (simple interface) molecular replacement program in the *Phenix* software package.[2] We used the structure of the Human Phospholipid Hydroperoxide Glutathione Peroxidase as a search model (PDB: 2OBI). Model building carried out using PHENIX and manual building using the program COOT.[3] Data collection and refinement statistics are summarized in **Supplementary Table S1**.

### Molecular Dynamics Simulations

#### GPx4–PIP complex structure preparation

The 3D structure of PIP-bound GPx4 complex was prepared by running multiple steered molecular dynamics (SMD) simulations. In these simulations, the PIP headgroup was directed toward identified protein hotspot residues by applying a harmonic bias to a defined collective variable (CV). This approach allows for the continuous evolution of the CV through an applied force, effectively modeling the binding process along a specific reaction coordinate. The hotspot residues that interact with the PIP headgroup were identified based on the co-crystallized structure of the GPx4–$IP_4$ complex. The minimum distances between the center-of-mass (COM) of $IP_4$ and the COMs of GPx4 residues were computed, then the six closest residues were selected as hotspot residues (**Supplementary Table S2**). The hotspot residues overlap with the residues identified as binding to $IP_4$ based on the co-crystallization result (**Figure 6**). Next, multiple SMD simulations were performed to pull the PIP headgroup close to the hotspot residues so that the prepared GPx4–PIP complex structure aligned well with the $IP_4$-bound GPx4 crystal structure (**Supplementary Figure S15**). All SMD steps were performed with harmonic potential using pull rate of 0.0005 nm/ps with harmonic constant of 1000 kJ $mol^{-1}$ $nm^{-2}$. After the completion of the SMD steps, the system was equilibrated under NPT conditions for 1 ns with position restraints on the protein backbone to relax the PIP structure, then a 50 ns unbiased simulation was performed as production. The production trajectories were analyzed to obtain time-series profiles of the distances between the PIP lipid termini and the COM of the GPx4 40s catalytic loop (**Supplementary Figure S16**). Specifically, distances were calculated between the COM of the GPx4 40s loop residues S44–E50 and the terminal carbon atoms of the saturated and unsaturated lipid tails of PIP.

**PIP hydroperoxides preparation**

To simulate the effects of lipid peroxidation on GPx4 membrane interactions, peroxidized PIP molecules were prepared using the PI(3,4,5)$P_3$ structure as a template. The initial PI(3,4,5)$P_3$ coordinates were obtained from CHARMM Small Molecule Library of CHARMM-GUI Archive

[4], and four peroxide groups were manually added to the unsaturated arachidonyl lipid tail (20:4) using Avogadro v1.2.0 (**Supplementary Figure S17**).[5] The resulting structure was used to generate the force field parameters and topology for the PIP hydroperoxides using the CHARMM-GUI *Ligand Reader & Modeler* module.[4,6] Following the approach described by Hu et al. 2025[7] to maintain consistency between the native and oxidized lipid species, all force field parameters for atoms outside the immediate peroxidation region were cross-referenced and reset to match the original PIP values. The residual charge discrepancy resulting from this alignment was evenly redistributed across the eight peroxide-region oxygens. This adjustment yielded partial charges that align with physically realistic values for oxygen atoms in aqueous or lipid environments, thereby maintaining the total molecular charge of -6.0 *e*.

**Membrane-bound GPx4 conformation preparation using SMD simulations**

To prepare membrane-bound GPx4 configurations, GPx4 was first positioned at ~1 nm above the membrane surface, then energy minimization and 1 ns equilibration were performed. We then performed multiple steps of unbiased MD and SMD to sample a membrane-bound conformation of GPx4 that was consistent with the binding pattern characterized by a previous MD study, where insertion of L130 was shown to be critical to the functional binding of GPx4 to the membrane.[7] All membrane systems were modeled using 1,2-dioleoyl-sn-glycero-3-phosphocholine (DOPC) lipids using the *Membrane Builder* module of CHARMM-GUI Input Generator.[4,8] All SMD simulations were performed using pull rate of 0.0005 nm/ps with harmonic constant of 1000 kJ $mol^{-1}$ $nm^{-2}$. All simulations of membrane systems were performed under NPT conditions using semi-isotropic barostat. Specific details for each membrane follow.

DOPC-only membrane system

After initial equilibration, temperature replica exchange MD (REMD) was performed to sample the membrane-bound conformation of GPx4.[9] REMD allows multiple replicas of a system to run in parallel at different temperatures, periodically exchanging configurations based on a Metropolis criterion to enhance sampling of conformational space. REMD for GPx4–DOPC membrane system was performed using 52 replicas with 2 K spacing between adjacent replicas, temperature starting from 298.15 to 400 K, for 50 ns allowing exchange every 1 ps.

While REMD simulations sampled membrane-bound states of GPx4, they failed to achieve the specific L130 insertion required, necessitating the use of SMD to force L130 insertion into the membrane. Therefore, a membrane-bound conformation with GPx4 closest to the membrane midplane from the trajectory of first replica at 298.15 K was selected to run the additional SMD to force L130 toward the midplane (**Supplementary Figure S18A**). L130 SMD was performed with same pull rate and harmonic constant introduced in the prior section while using the cylindrical pull coordinate geometry implemented in GROMACS. The *z*-component of the distance between the COM of the bilayer and the COM of the pulled L130, *dz*, was used as the CV for L130 SMD. By using the cylindrical pull coordinate geometry, only the portion of the bilayer within a cylindrical region around the pulled loop was used when computing the COM of the bilayer, which is accomplished by weighting the contribution of each atom to the calculation of the reaction coordinate by a factor $w_i$ which is defined in Equation S1:

$$w_i = \begin{cases} 1 - 2\left(\frac{r_i}{r_{cyl}}\right)^2 + \left(\frac{r_i}{r_{cyl}}\right)^4, & r_i < r_{cyl} \\ 0, & r_i \geq r_{cyl} \end{cases} \tag{S1}$$

Here, $r_i$ is the radial distance between atom $i$ and the COM of the pulled L130 in the $x$-$y$ plane and $r_{cyl} = 1.5$ nm for all L130 SMDs. After the L130 insertion, unbiased MD was performed for 100 ns to sample GPx4–membrane binding patterns.

PIP-containing DOPC membrane system

Unlike the DOPC-only membrane system, additional steps were performed to ensure initial GPx4-PIP binding for the membranes containing PIP. After initial equilibration, SMD was performed using the distance between the COM of PIP headgroup and GPx4 hotspot residues as the CV (**Supplementary Figure S18B**), thereby following a similar procedure as was used to prepare the GPx4-PIP complex in bulk solution. Next, L130 SMD was performed using same conditions introduced in the prior section, resulting in a functional membrane-bound GPx4 structure with hotspot residues interacting with PIP headgroup in a similar manner to $IP_4$ binding and with L130 inserted into the membrane. From this configuration, a 100 ns unbiased MD was performed as production to sample GPx4-membrane binding patterns. The same procedure was used to prepare the PIP hydroperoxide-containing DOPC membrane systems.

**Contact barcode plot**

The final 10 ns of unbiased MD trajectories for which GPx4 and membrane are in contact provide reliable information to investigate protein-membrane binding patterns.[10] Therefore, the last 10 ns of each unbiased MD trajectory of GPx4–membrane systems was used to investigate composition-dependent binding patterns (**Supplementary Figure S19**). For each simulation configuration, protein residues with any heavy atom within 0.5 nm of any heavy atom of a lipid on the membrane surface were designated as in contact. **Supplementary Figure S19** shows the residues in contact (black) as a function of simulation time for all trajectories. **Figure 7C** of the main text shows the time-averaged fraction of the simulation trajectory in which each residue is in contact with the membrane, which is designated as the contact fraction.

**Umbrella Sampling**

Umbrella sampling (US) is a method for calculating a free energy profile as a function of a predefined reaction coordinate.[11] A harmonic biasing potential, $V_\xi$, is applied to enhance sampling around specific points, $\xi_0$, of the reaction coordinate:

$$V_\xi = \frac{1}{2}k(\xi - \xi_0)^2 \tag{S2}$$

A series of independent simulations, referred to as windows, are performed using different values of $\xi_0$ to sample configurations across the full range of the reaction coordinate. When neighboring windows provide sufficient configurational overlap, methods such as the Weighted Histogram Analysis Method (WHAM)[12] can be used to combine the biased simulations and reconstruct the potential of mean force (PMF), corresponding to the free energy as a function of the reaction coordinate.

In this study, US simulations were performed to compute PMFs for protrusion of the unsaturated lipid tail toward the GPx4 40s loop. The temperature was maintained at 298.15 K using a velocity-rescale thermostat with a time constant of 1.0 ps and the pressure was maintained at 1 bar using a semi-isotropic stochastic cell-rescale barostat with a time constant of 5.0 ps and a compressibility of 4.5 x $10^{-5}$ $bar^{-1}$. For each PIP-containing system, 42 or 24 umbrella windows were used, with adjacent windows spaced by 0.1 nm along the reaction coordinate (**Supplementary Table S3**). The reaction coordinate was defined as the distance between the COM of the unsaturated lipid tail terminal groups and the COM of the GPx4 40s loop, computed using the distance pull coordinate geometry. Initial configurations for the windows were selected from SMD trajectories in which the unsaturated lipid tail was pulled toward the COM of the 40s loop across the bilayer. These SMD simulations were initiated from the final snapshot of a 100 ns unbiased MD simulation of membrane-bound GPx4 across replicas (**Supplementary Figure S19B-C**).

Each window was simulated with a harmonic force constant of 1000 kJ $mol^{-1}$ $nm^{-2}$ to restrain sampling around the target reaction coordinate value. For both the PIP- and PIP-hydroperoxide-containing DOPC membrane systems, each window was simulated for 100 ns. Simulation lengths were selected based on convergence of the resulting PMF profiles and corresponding histograms (**Supplementary Figs. S22-S25**). Depending on the system and convergence behavior, the initial 10 to 50 ns of the US trajectory were discarded as equilibration, and the remaining trajectory frames were used for PMF reconstruction.

PMFs were computed using WHAM, and uncertainties were estimated by bootstrapping.[12] WHAM was performed using 200 bins along the reaction coordinate. Bootstrapping was performed 200 times using the histograms generated from the umbrella windows (**Supplementary Figs. S23, S25**). For each window, the integrated autocorrelation time, $\tau$, was estimated and used to weight the corresponding histogram by $\frac{1}{1+2\tau/\mathrm{d}t}$.

**Convergence of Potentials of Mean Force**

To assess convergence of the PMFs calculated from US for the PIP unsaturated lipid tail and the GPx4 40s loop, each US trajectory was systematically split into distinct "equilibration" and "production" increments.[13] For each split, the initial portion of the trajectory corresponding to the equilibration time was discarded, and the remaining trajectory frames, corresponding to the production time, were used to compute a PMF using WHAM.

Each PMF was plotted as a function of the distance between the COM of the pulled PIP unsaturated lipid tail and the COM of the GPx4 40s loop, $d_{COM}$. The unsaturated lipid tail was in contact with the GPx4 40s loop at $d_{COM} = 0.5$ nm. In contrast, the lipid tail was in its most equilibrated bilayer-embedded state at the maximum $d_{COM}$ value sampled for each PIP-containing system (**Supplementary Figs. S22A, S24A**). Different equilibration/production time intervals were evaluated by calculating the root mean squared error (RMSE) between PMFs generated using consecutive equilibration and production intervals. Before comparison, the global minimum of each PMF was shifted to zero to ensure consistent alignment. The equilibration/production interval with the lowest RMSE relative to the preceding and following intervals, indicated in bold in the tables of **Supplementary Figs. S22B and S24B**, was selected as the converged interval and used to generate the PMFs shown in the main text.

For **Supplementary Figs. S22B and S24B**, the legends of the replica 1 PMF plots list the equilibration/production intervals tested for each system. In the legends, “eq” denotes equilibration time and “prd” denotes production time (e.g., “eq10prd90” indicates the first 10 ns of a 100 ns simulation are discarded as equilibration time while the remaining 90 ns are considered as production time and used in WHAM to calculate the PMF). The shaded area indicates the standard deviation estimated by bootstrapping. The tables next to the PMF convergence plots report RMSE values between PMFs computed with consecutive sets of time increments. $\Delta G_{protrusion}$ is defined as the highest free energy barrier that the unsaturated lipid tail has to overcome during the protrusion process.

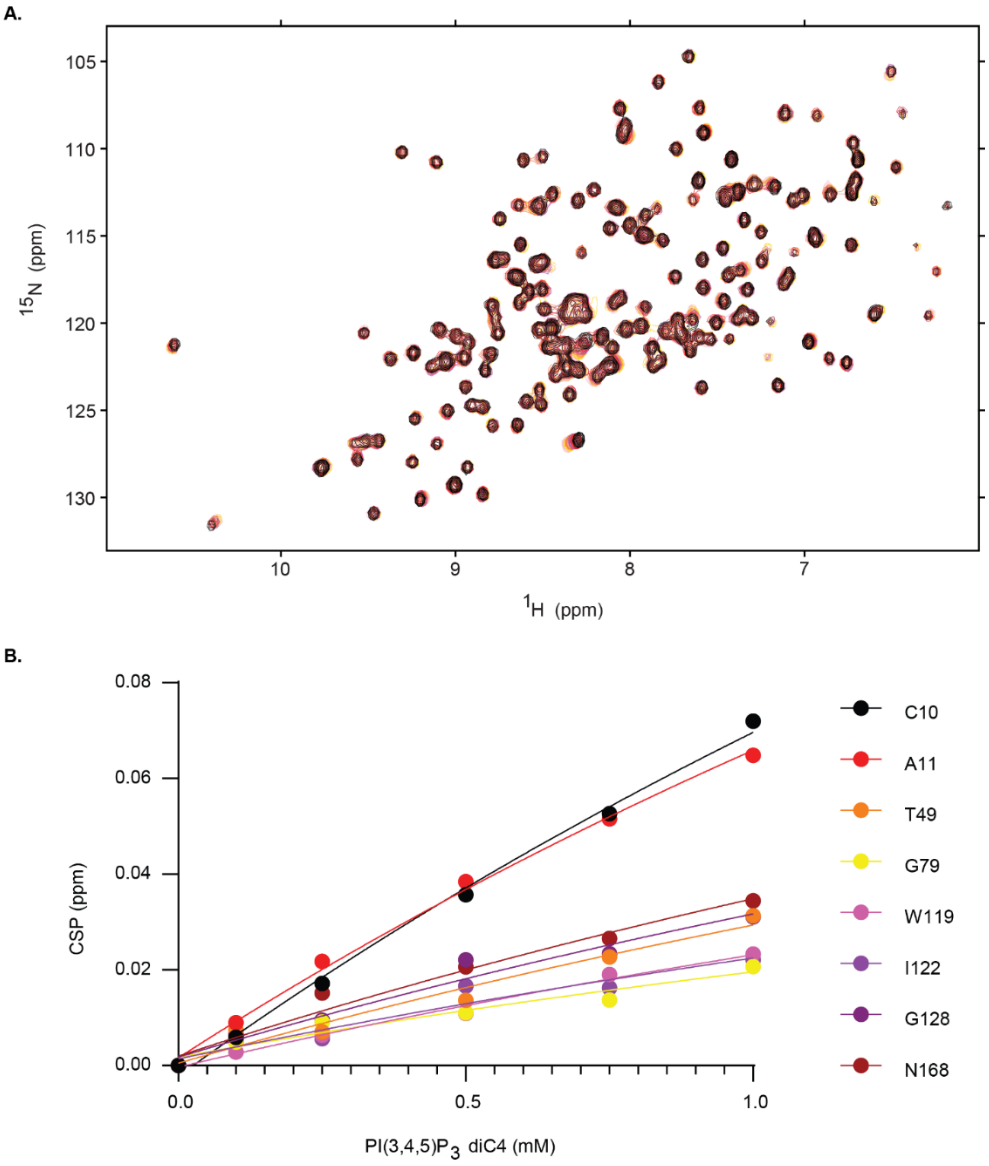


**Supplementary Figure S1.** Titration of PI(3,4,5)$P_3$ diC4 with 124 µM GPx4 observed with $^{1}H$-$^{15}N$ HSQCs allows estimation of an apparent $K_d$ = 4.6 ± 2.1 mM, though lack of saturation makes an accurate determination difficult. **A.** $^{15}N$-HSQC overlay of apo GPx4 (pink) titrated with PI(3,4,5)$P_3$ diC4 in the following sequence: 1 mM (black), 0.75 mM (red), 0.5 mM (magenta), 0.25 mM (coral), 0.1 mM (gold). **B.** Plot of representative CSPs used to extract an apparent $K_d$ value.

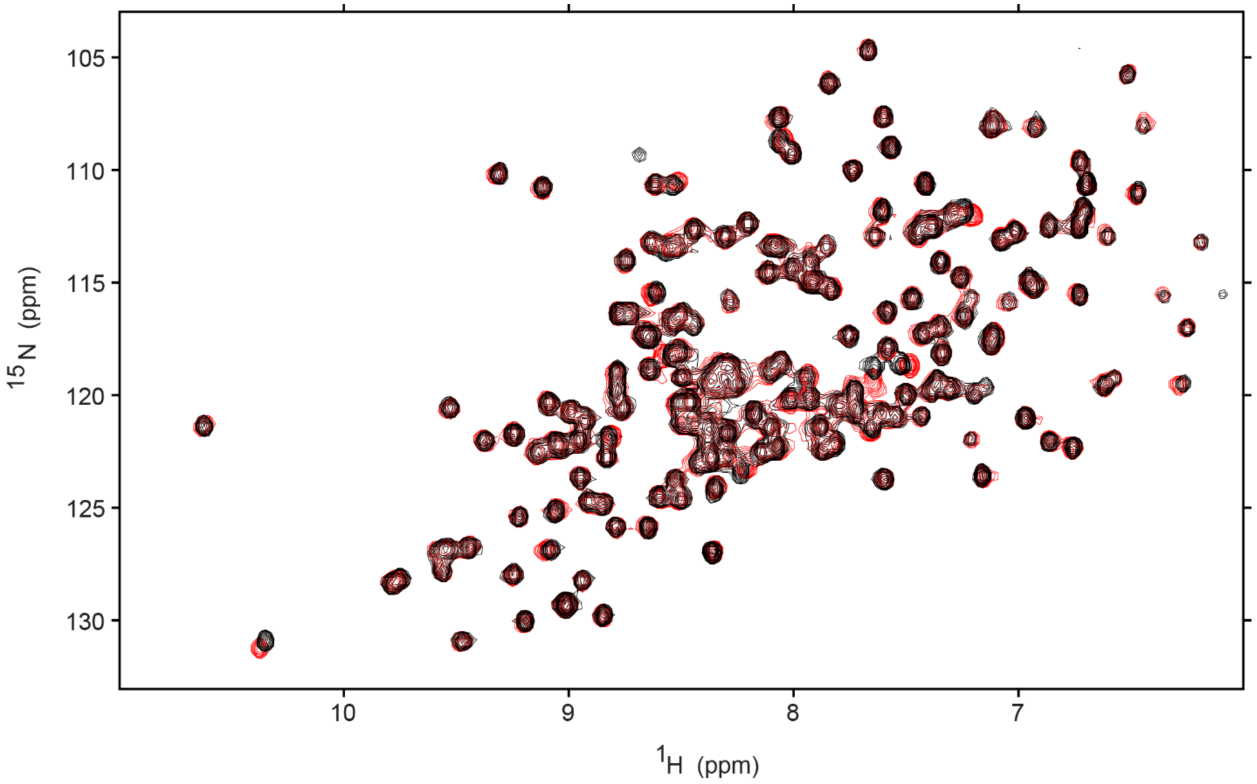


**Supplementary Figure S2.** $^{15}$N-HSQC overlay of apo 124 µM GPx4 (red) with 124 µM GPx4 plus 2 mM o-phosphor-L-serine sodium salt hydrate (black).

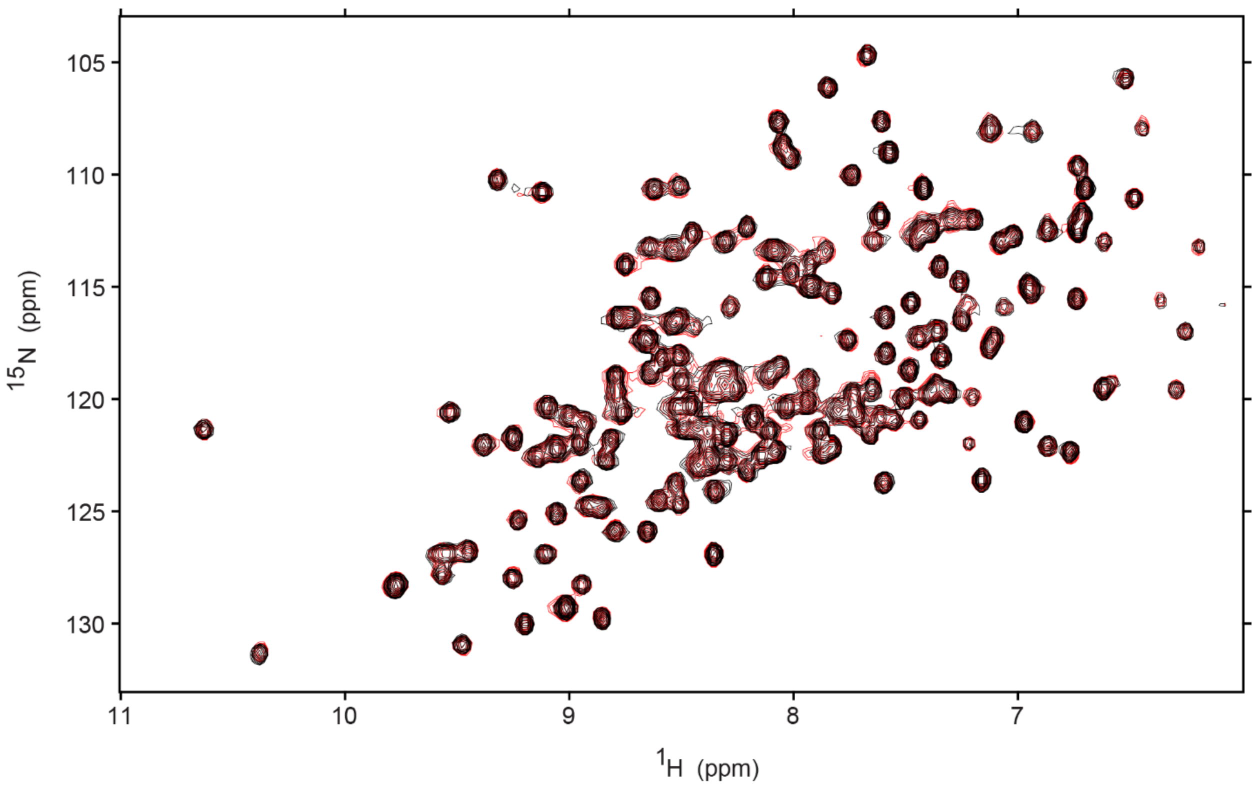


**Supplementary Figure S3.** $^{15}$N-HSQC overlay of apo 124 μM GPx4 (red) with 124 μM GPx4 plus 2 mM rac-glycerol 1-phosphate sodium salt hydrate (black).

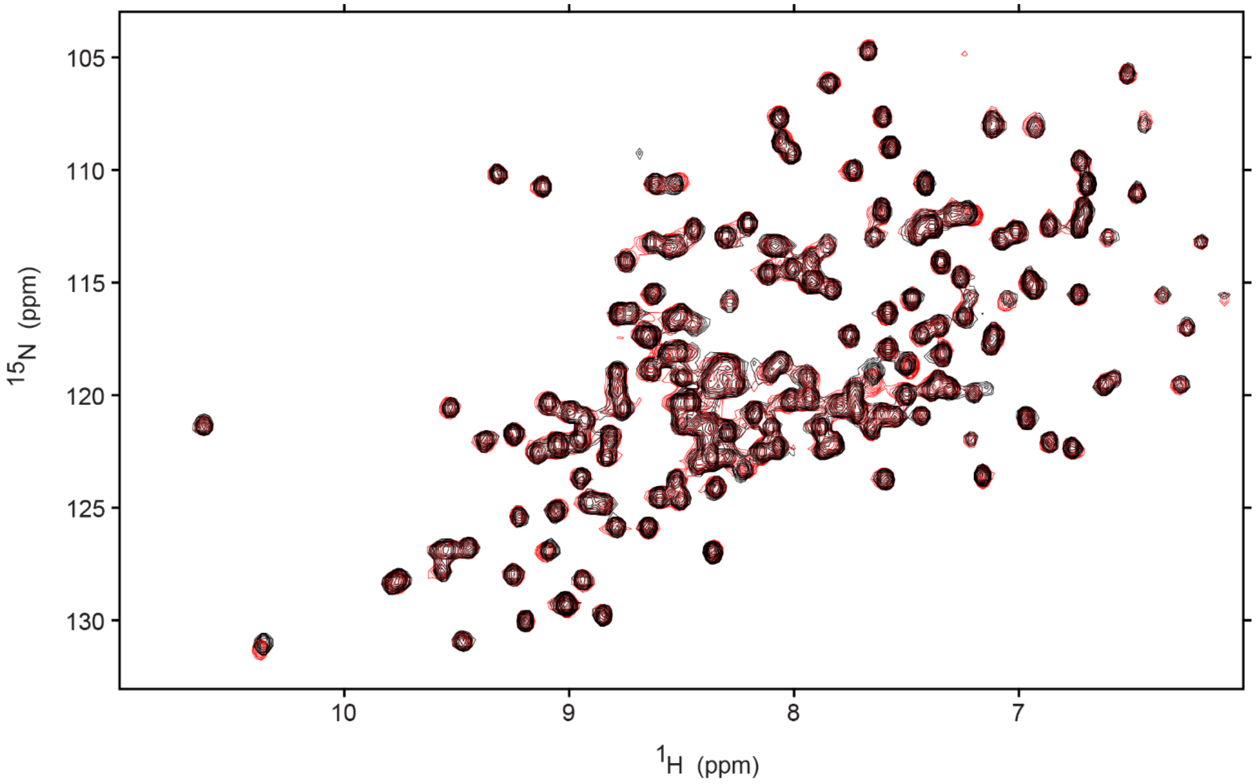


**Supplementary Figure S4.** $^{15}$N-HSQC overlay of apo 124 µM GPx4 (red) with 124 µM GPx4 plus 2 mM o-phosphoethanolamine (black).

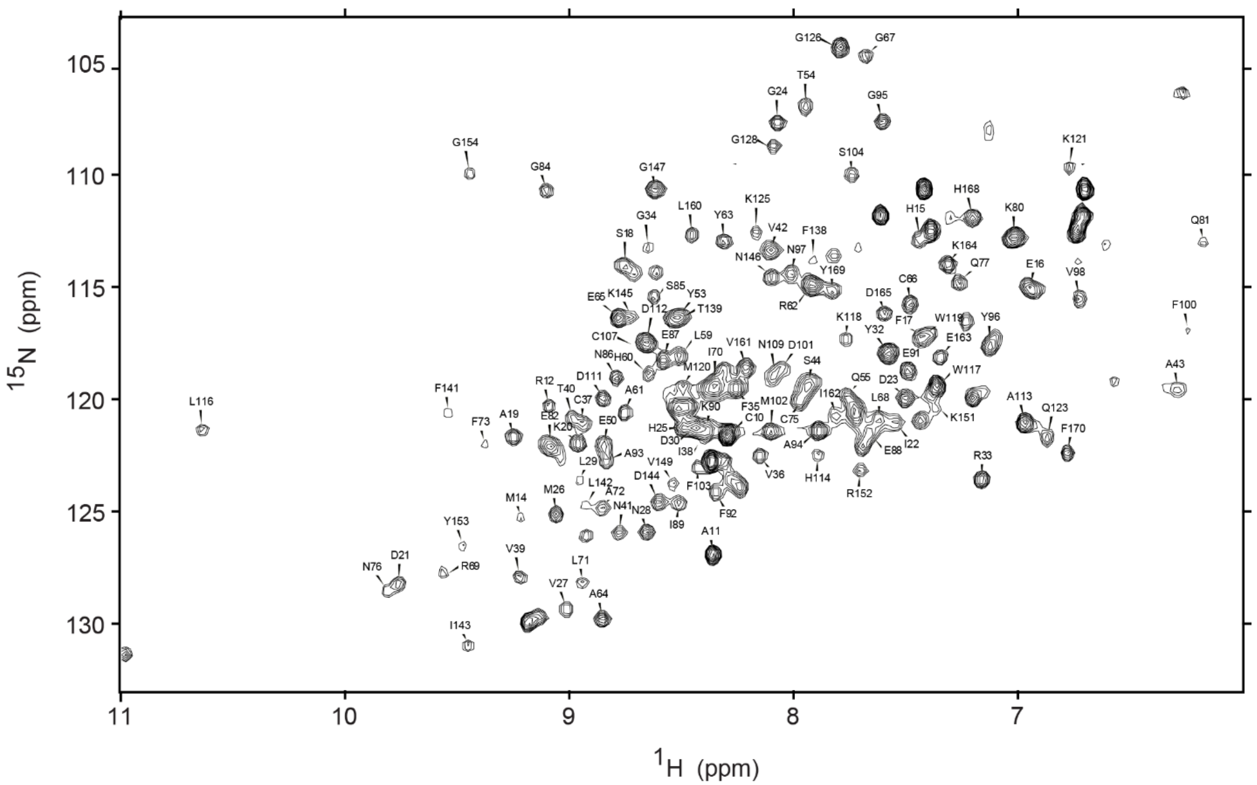


**Supplementary Figure S5.** NMR residue assignments of U46C-GPx4 in 25 mM DPC micelles. $^{1}H$-$^{15}N$ HSQC of 375 μM protein collected at 25 °C and 700 MHz in 20 mM Bis-Tris pH 6.0, 100 mM NaCl, and 10 mM DTT with 10% (v/v) $D_2O$. Peaks are labeled with the corresponding amino acid assignment.

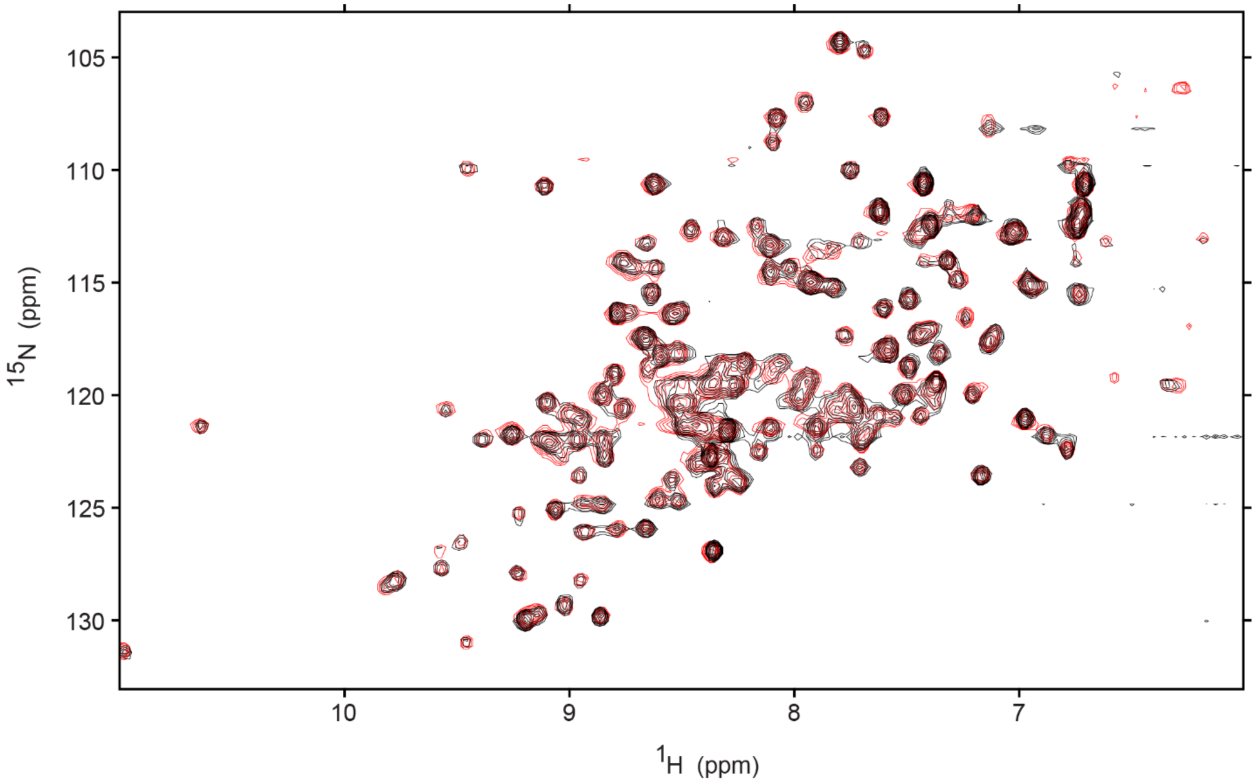


**Supplementary Figure S6.** $^{15}$N-HSQC overlay of 124 µM GPx4 in 25 mM DPC micelles (red) with the addition of 500 µM 1,2-dilauroyl-sn-glycero-3-phosphoethanolamine (PE diC12) (black) showing minimal CSPs.

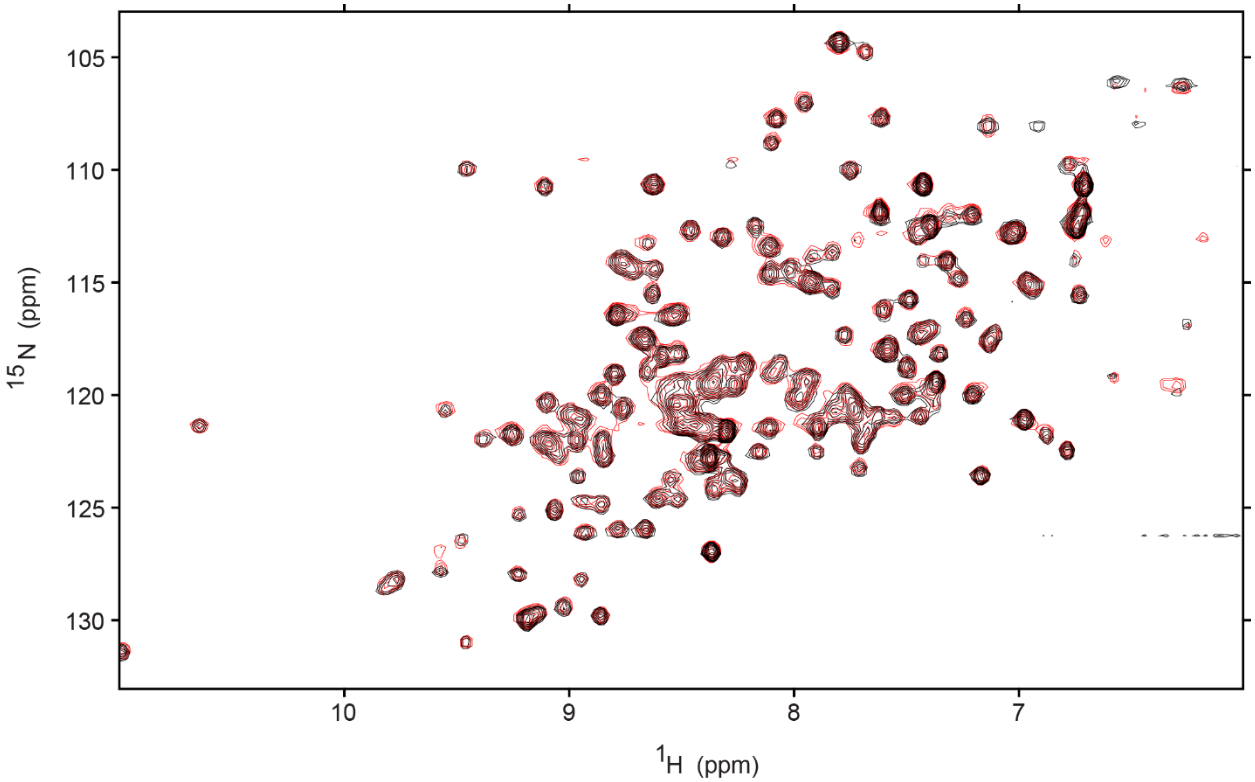


**Supplementary Figure S7.** $^{15}$N-HSQC overlay of 129 μM GPx4 in 25 mM DPC micelles (red) with the addition of 500 μM 1,2-dipalmitoyl-sn-glycero-3-phosphoserine (16:0 PS) (black) with minimal CSPs.

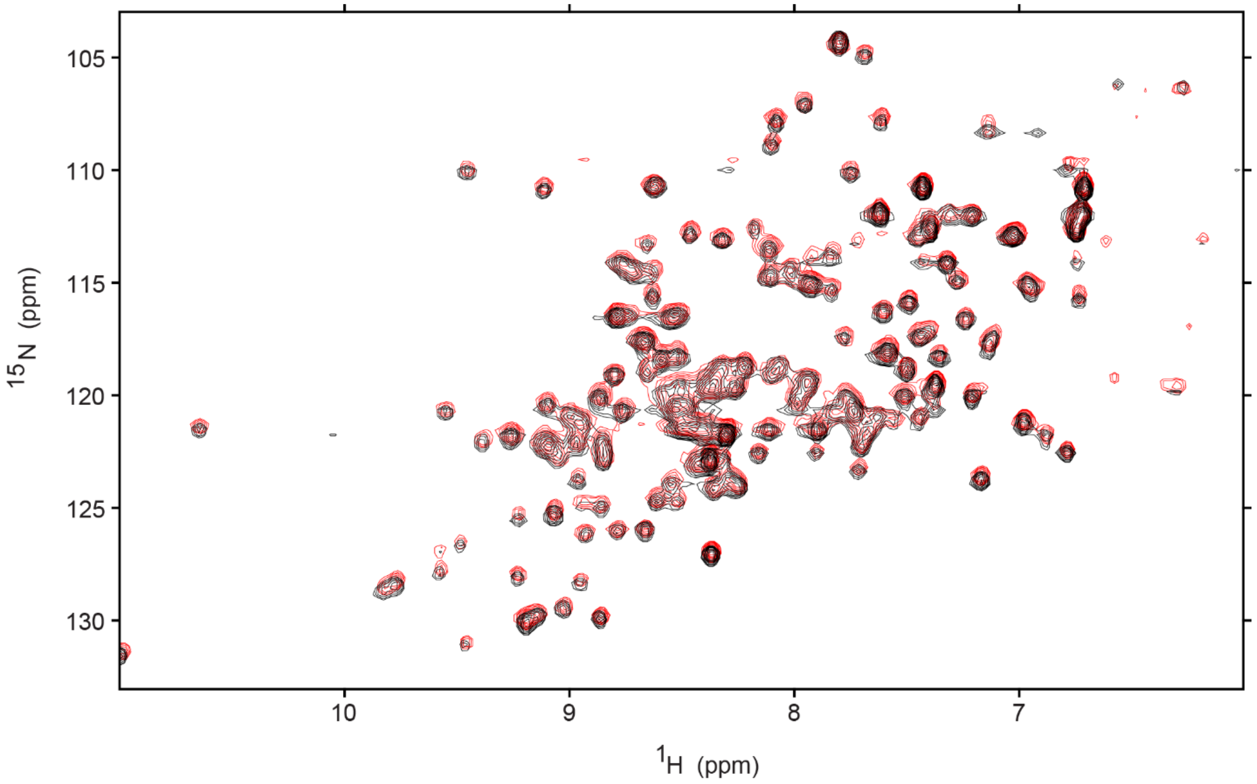


**Supplementary Figure S8.** $^{15}$N-HSQC overlay of 129 µM GPx4 in 25 mM DPC micelles (red) with the addition of 500 µM 1,2-dilauroyl-sn-glycero-3-phosphate (12:0 PA) (black) with minimal CSPs.

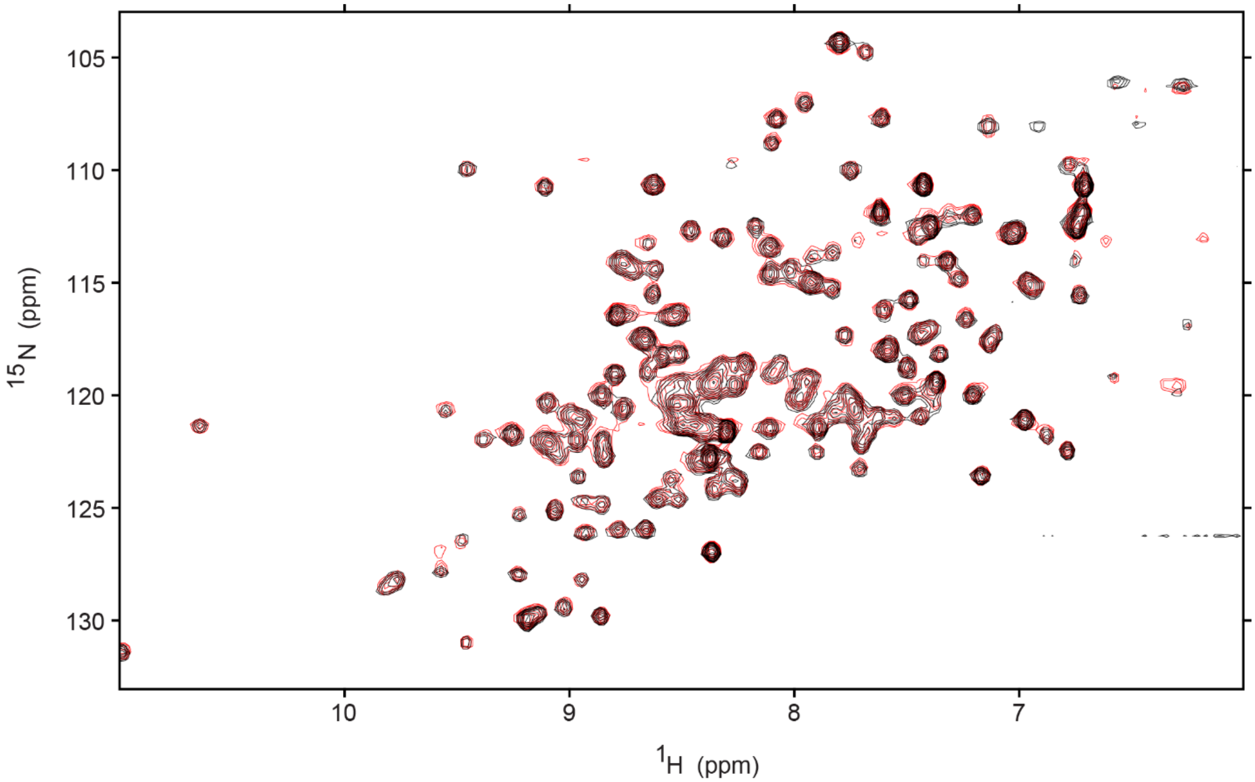


**Supplementary Figure S9.** $^{15}$N-HSQC overlay of 129 μM GPx4 in 25 mM DPC micelles (red) with the addition of 500 μM 1,2-dilauroyl-sn-glycero-3-phospho-(1'-rac-glycerol) (12:0 PG) (black) with minimal CSPs.

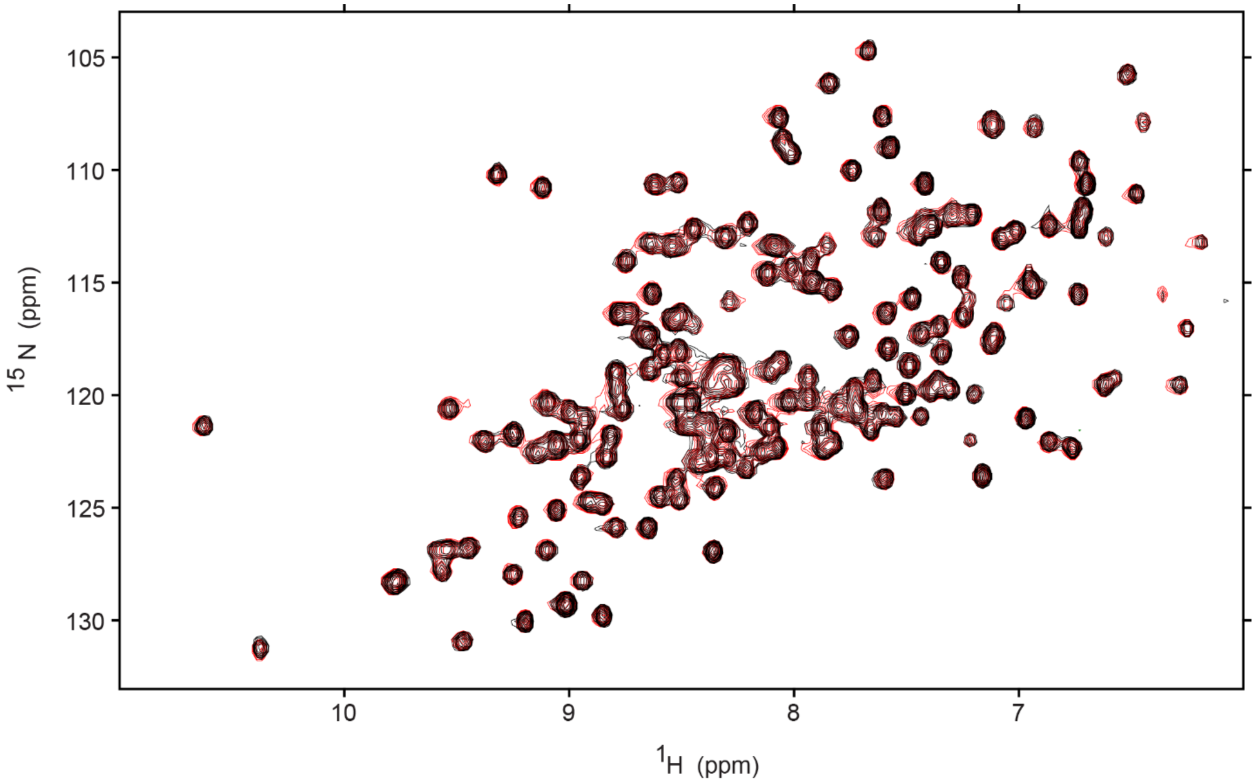


**Supplementary Figure S10.** $^{15}$N-HSQC overlay of apo 124 µM GPx4 (red) with the addition of 2 mM $IP_1$ (1) (black).

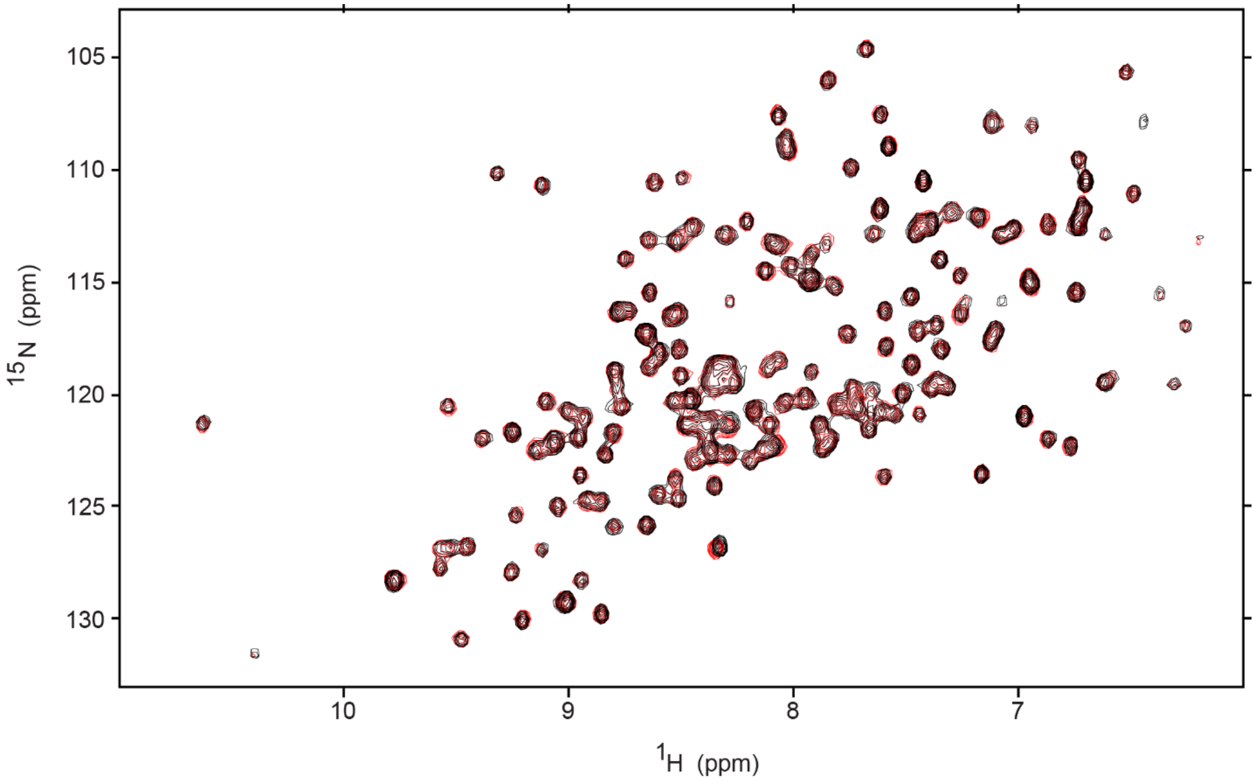


**Supplementary Figure S11.** $^{15}$N-HSQC overlay of apo 124 µM GPx4 (red) with the addition of 1 mM $IP_3$ (1,3,4) (black).

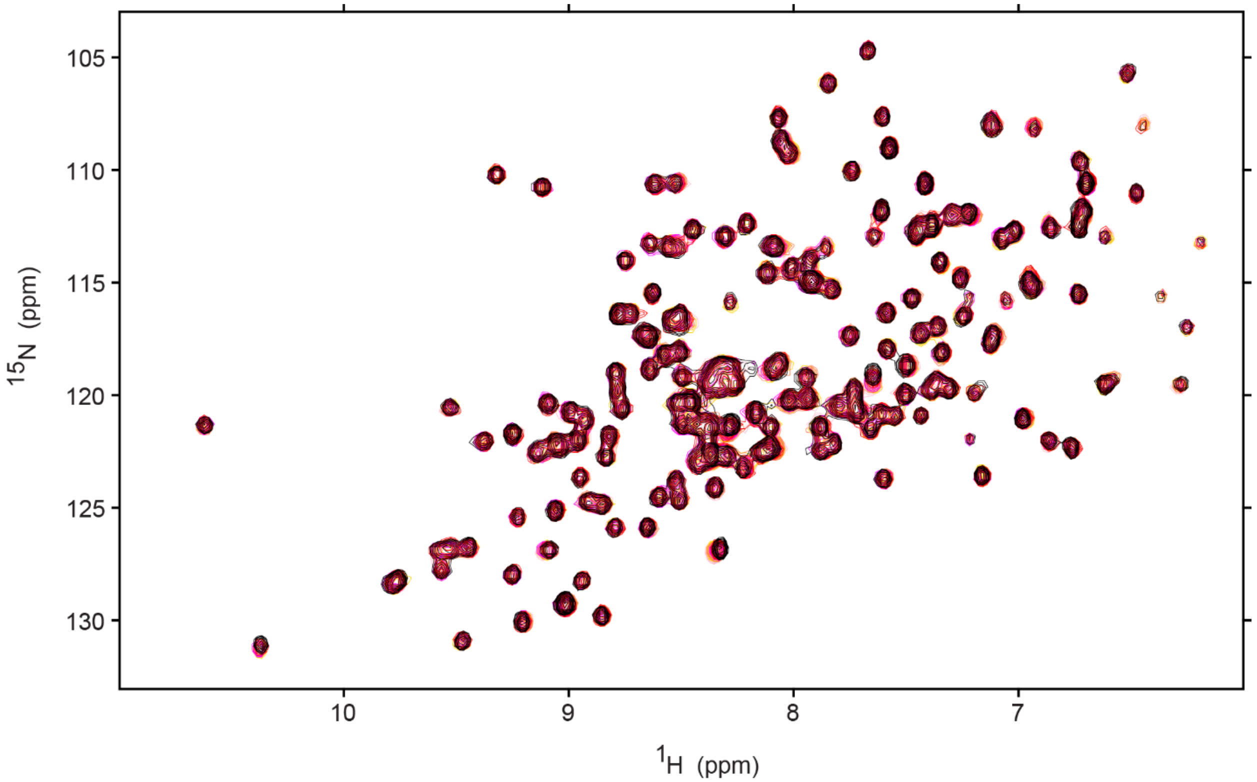


**Supplementary Figure S12.** $^{15}$N-HSQC overlay of apo GPx4 (pink) titrated with $IP_3$ (1,3,5) in the following sequence: 1 mM $IP_3$ (black), 0.75 mM $IP_3$ (red), 0.5 mM $IP_3$ (magenta), 0.25 mM $IP_3$ (coral), 0.1 mM $IP_3$ (gold).

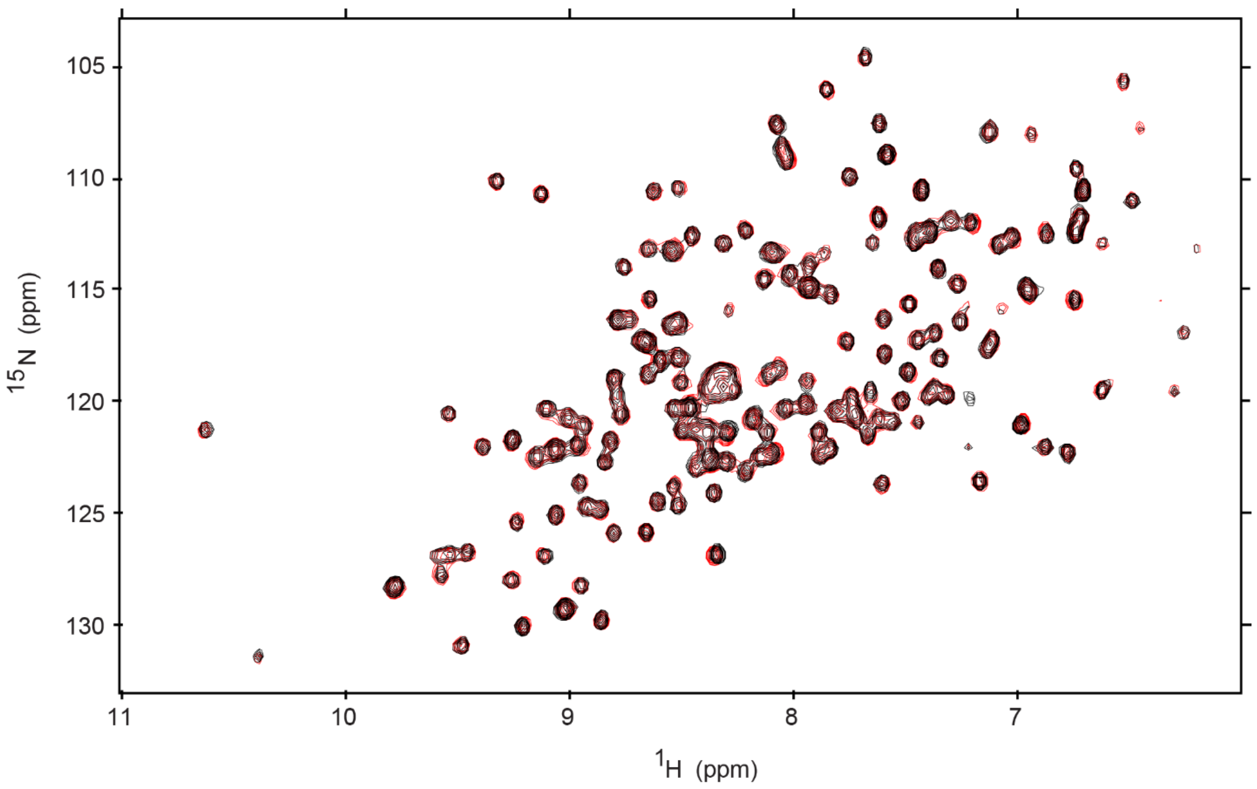


**Supplementary Figure S13.** $^{15}$N-HSQC overlay of apo 124 μM GPx4 (red) with the addition of 1 mM $IP_3$ (1,4,5) (black).

**Supplementary Table S1**: Data Collection and refinement statistics for PDB ID: 35WP

| **Data Collection Statistics** | |
|---|---|
| Space group | P 2 1 |
| Unit-cell *a, b, c* (Å) | 76.906, 61.884, 80.803, β = 113.40 (°) |
| Resolution (Å)<br>Total reflections | 26.90–2.50 (2.60 – 2.50)<br>233198 (26115) |
| Unique reflections | 24371 (2767) |
| Redundancy | 9.6(9.4) |
| Completeness (%) | 99.9 (100.0) |
| Average **I**/σ(I) | 16.6 (2.6) |
| $R_{merge}$ (%)[a] | 11.4 (93.6) |
| **Refinement Statistics** | |
| Resolution (Å) | 26.90-2.50 (2.59-2.50) |
| No. of reflections | 24288 (2437) |
| $R_{work}$ (%) | 23.00 (28.74) |
| $R_{free}$ (%)[b] | 29.58 (36.37) |
| R.m.s.d. bonds (Å) | 0.008 |
| R.m.s.d. angles (°) | 0.990 |
| Dihedral angles | |
| Most favored (%) | 95.05 |
| Allowed (%) | 4.31 |
| **Average B (Å$^2$) / atoms** | |
| All atoms | 44.75 |
| Protein<br>Ligand<br>Solvent<br>**Number of non-hydrogen atoms** | 44.18<br>104.87<br>40.22<br>5221 |
| Macromolecules<br>Ligands<br>Water | 5065<br>56<br>100 |
| Protein residues | 643 |

[a]$R_{merge} = \Sigma_{hkl}\Sigma_i|I_i(hkl) - <I(hkl)>|/\Sigma_{hkl}\Sigma_i I_i(hkl)$. [b]$R_{free}$ was calculated from 5% randomly selected reflection for cross-validation. All other measured reflections were used during refinement.

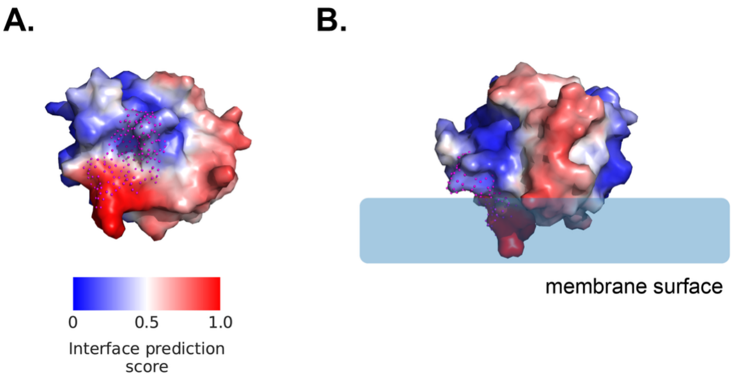


**Supplementary Figure S14.** Surface-level membrane-binding site predictions for GPx4 using MaSIF-PMP. **A.** Snapshots show MaSIF-PMP prediction scores mapped onto the surface of GPx4 (PDB ID: 2OBI). Prediction scores range from 0, indicating nonbinding regions, to 1, indicating predicted membrane-binding regions, and are visualized using a blue-to-red color scale. Surface representations are aligned with the GPx4 ribbon representations shown in **Figure 6A** and **6B** of the main text. Magenta spheres indicate surface regions corresponding to $IP_4$-binding residues (N109–D111, K121, and G131–A133) shown in **Figure 6**. **B.** The expected membrane surface was assigned based on the location of the highest-scoring predicted surface regions, which were centered around L130 and adjacent residues. Overall, MaSIF-PMP predictions identify membrane-binding regions on the GPx4 surface that overlap well with the $IP_4$-binding site.

**Supplementary Table S2.** Hotspot residues closest to $IP_4$ and corresponding center-of-mass distances ($d_{COM}$).

| Residues | $d_{COM}$ (nm) |
|---|---|
| G110 | 0.6235 |
| A133 | 0.6793 |
| N109 | 0.7017 |
| N132 | 0.7029 |
| K121 | 0.7386 |
| K118 | 0.8017 |

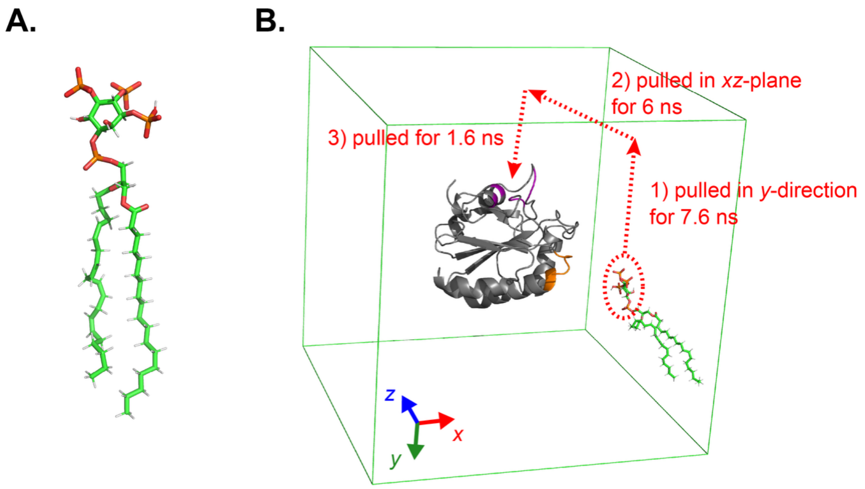


**Supplementary Figure S15.** GPx4–PIP complex structure preparation. **A.** Atomistic model of PI(3,4,5)$P_3$ (18:0/20:4), which was used as the native PIP model. PIP is shown in a stick representation with carbon atoms in green, hydrogen atoms in white, oxygen atoms in red, and phosphorus atoms in orange. **B.** Multiple SMD simulations to prepare a GPx4–PIP complex structure that aligned well with the $IP_4$-bound GPx4 co-crystal structure. GPx4 is shown in a gray ribbon representation, with the 40s loop highlighted in orange and hotspot residues in magenta. The color scheme of PIP follows the same definition used in **A.** Water molecules and ions are not shown to aid visualization. The periodic boundary of the system is indicated by a green box. The collective variable for SMD was defined as the distance between the center-of-mass (COM) of the PIP headgroup and the COM of the six hotspot residues. Red arrows and annotations denote the pulling direction, orientational bias, and simulation duration (ns) for each stage.

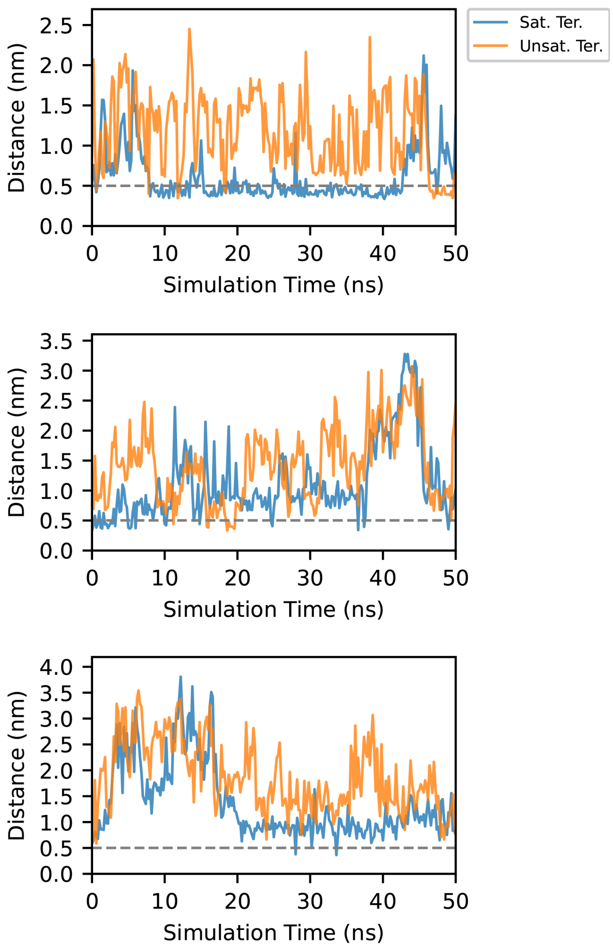


| | Sat. Ter. | Unsat. Ter. |
|---|---|---|
| Replica 1 | 0.611 ± 0.320 nm | 1.167 ± 0.473 nm |
| Replica 2 | 1.125 ± 0.639 nm | 1.419 ± 0.610 nm |
| Replica 3 | 1.371 ± 0.721 nm | 1.872 ± 0.667 nm |

**Supplementary Figure S16.** Time-series distance profiles between PIP lipid termini and the GPx4 40s catalytic loop from three independent simulation replicas in bulk water. Distances were calculated between the center-of-mass of the 40s loop and either the saturated lipid terminus (Sat. Ter.) or the unsaturated lipid terminus (Unsat. Ter.) for each replica. A contact threshold of 0.5 nm is indicated by the gray dashed line. The accompanying table provides the corresponding averages and standard deviations of the distance distributions for both the saturated and unsaturated termini throughout the simulation trajectories.

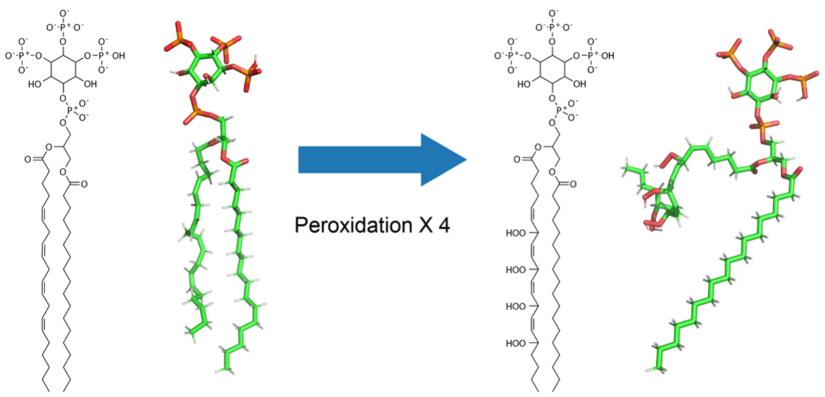


**Supplementary Figure S17.** Chemical structures and atomistic models of native and peroxidized PIP. The native PI(3,4,5)$P_3$ (18:0/20:4) structure served as the template for the peroxidized model. Four hydroperoxide groups were manually incorporated into the unsaturated arachidonyl lipid tail of the template structure to generate the peroxidized species. The color scheme of PIP species follows the same definition used in **Supplementary Figure S15A**.

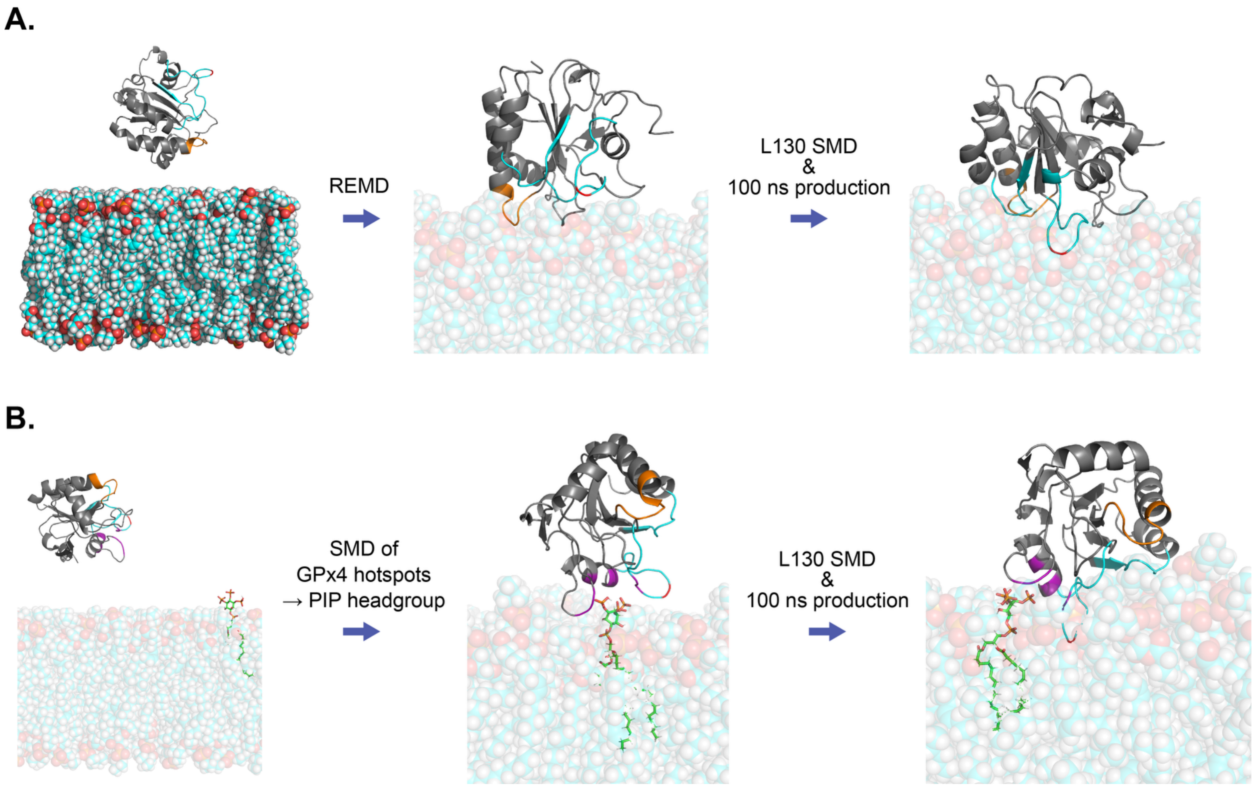


**Supplementary Figure S18.** Membrane-bound GPx4 conformation preparation. **A.** DOPC-only membrane system. The color scheme of GPx4 follows the same definition used in **Supplementary Figure S15B**, with the cationic regions highlighted in cyan and L130 in red. DOPC lipids are shown in van der Waals representation using the same color scheme as PIP lipids, except that carbon atoms are colored cyan. Water molecules and ions are not shown to aid visualization. **B.** PIP-containing DOPC membrane system. Representative snapshots of intermediate configurations of the preparation procedure for preparing native-PIP-containing membrane system are shown as an example. The same procedure was used for the PIP-hydroperoxide-containing membrane systems. The color scheme follows the same definition used in **A.** and **Supplementary Figure S15A**.

A.

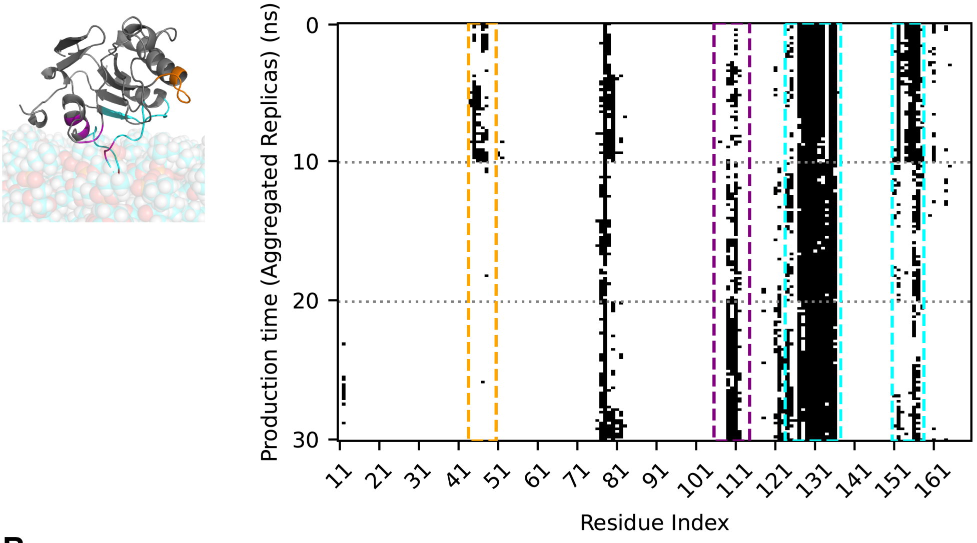

Production time (Aggregated Replicas) (ns)
0
10
20
30
11 21 31 41 51 61 71 81 91 101 111 121 131 141 151 161
Residue Index


B.

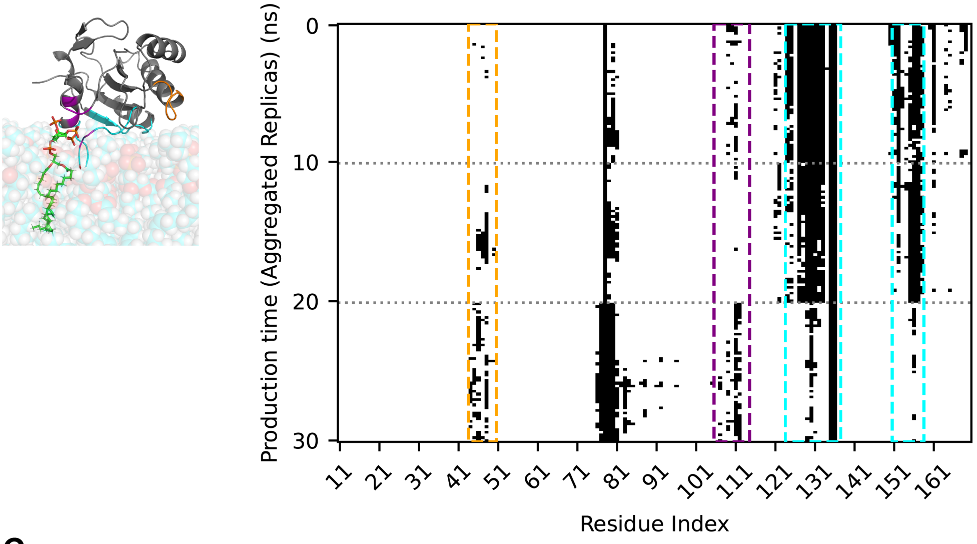

Production time (Aggregated Replicas) (ns)
0
10
20
30
11 21 31 41 51 61 71 81 91 101 111 121 131 141 151 161
Residue Index


C.

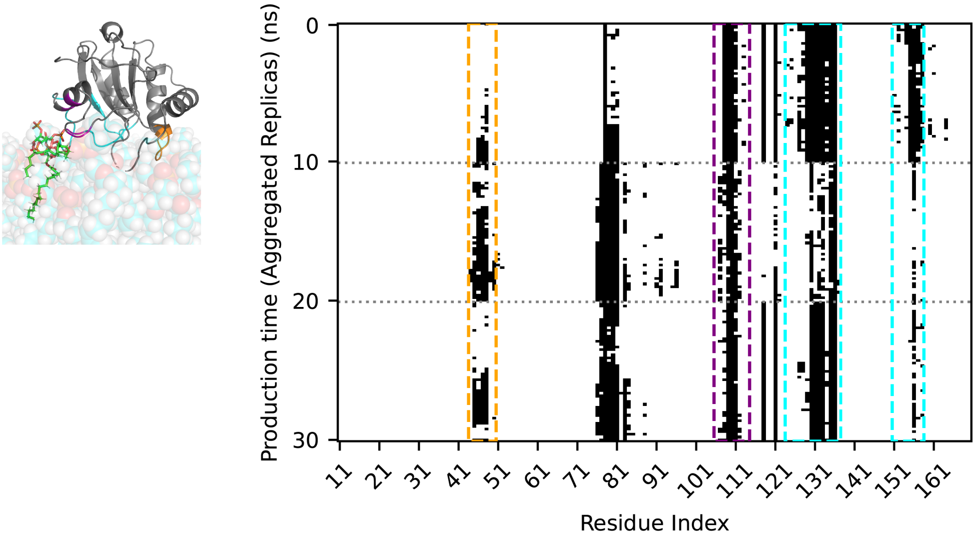

Production time (Aggregated Replicas) (ns)
0
10
20
30
11 21 31 41 51 61 71 81 91 101 111 121 131 141 151 161
Residue Index

**Supplementary Figure S19.** Contact barcode plots and representative snapshots of membrane-bound GPx4 across three lipid environments. Results are shown for (**A**) DOPC-only, (**B**) DOPC + PIP, and (**C**) DOPC + PIP-hydroperoxide membranes, with a representative snapshot of membrane-bound GPx4 provided for each. Data were aggregated across three replicas per system, with individual replica boundaries delimited by gray dashed lines. The color scheme follows the same definition used in **Supplementary Figure S18**. In the barcode plots, black and white indicate the presence or absence of contact between protein residues and membrane lipids, respectively. Dashed boxes highlight key functional regions: orange for the 40s catalytic loop, magenta for areas proximal to hotspot residues, and cyan for cationic regions. Two catalytic triad residues, U46 and W136, are located within the 40s loop and cationic regions, respectively, while Q81 is clearly identifiable within the barcode plot.

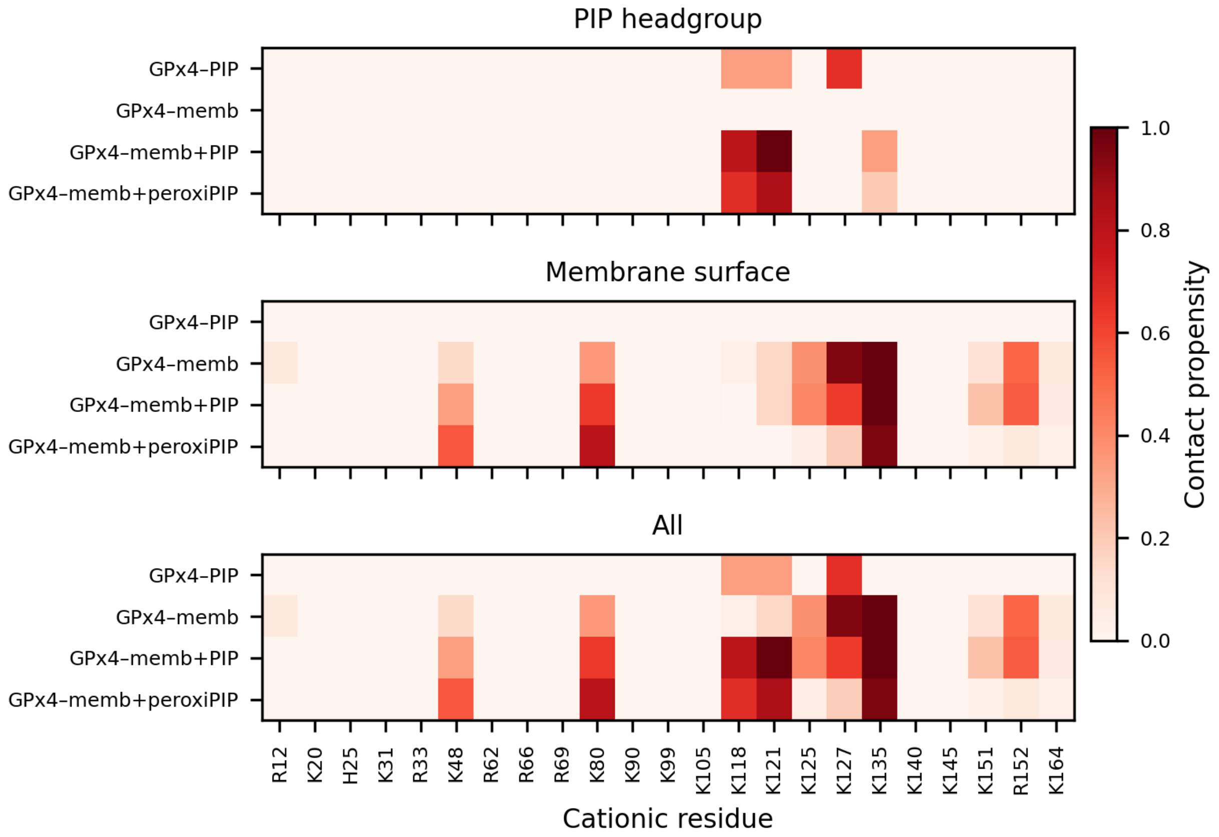


**Supplementary Figure S20.** Heatmap of contact propensities of GPx4 cationic residues across simulation systems and replicates. Contact propensities were computed as the fraction of production frames in which each residue was in contact with PIP headgroups, other membrane lipids, or all lipids combined, using the production trajectories from all replicates. "GPx4–PIP" represents the aqueous GPx4–PIP complex, while "memb," "memb+PIP," and "memb+peroxiPIP" represent DOPC membrane systems without PIP, with native PIP, and with peroxidized PIP, respectively.

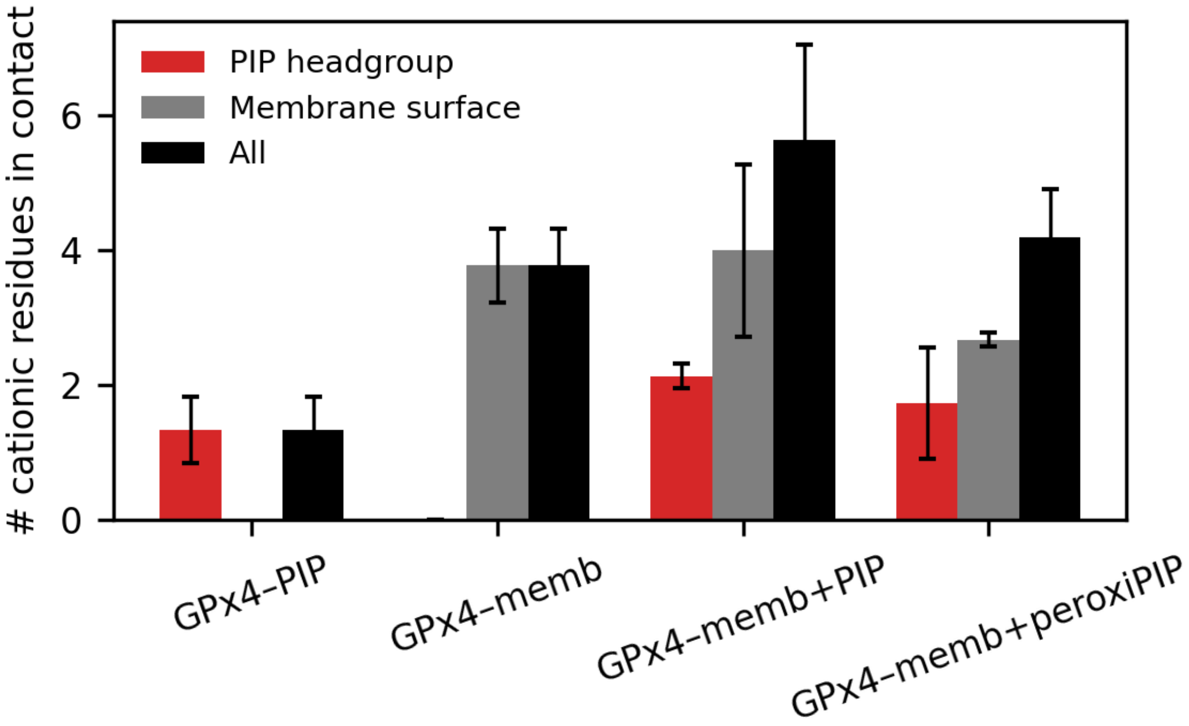


**Supplementary Figure S21.** Number of cationic residues in contact with the membrane. Labels follow the definitions provided in **Supplementary Figure S20**. Mean values and standard deviations were computed across replicate simulations for each system.

**Supplementary Table S3.** Number of components and umbrella sampling (US) windows for each simulated system.

| | GPx4 molecule | DOPC molecules | PIP molecule | Water molecules | $Na^{+}$ ions | $Cl^{-}$ ions | Total atoms | # of US windows |
|---|---|---|---|---|---|---|---|---|
| GPx4–PIP | 1 | 0 | 1 | 16,504 | 35 | 31 | 52,301 | N/A |
| GPx4–DOPC membrane | 1 | 160 | 0 | 15,691 | 27 | 29 | 71,841 | N/A |
| GPx4–(DOPC+PIP) membrane | 1 | 300 | 1 | 30,696 | 58 | 54 | 136,323 | 42 |
| GPx4–(DOPC+PIP hydroperoxide) membrane | 1 | 300 | 1 | 30,544 | 60 | 56 | 135,879 | 24 |

**A.**

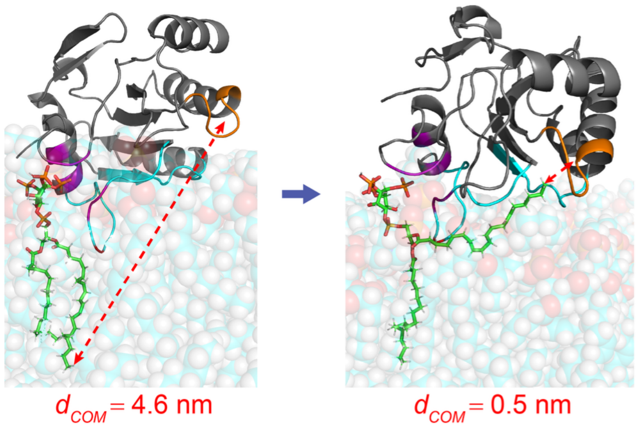


**B.**

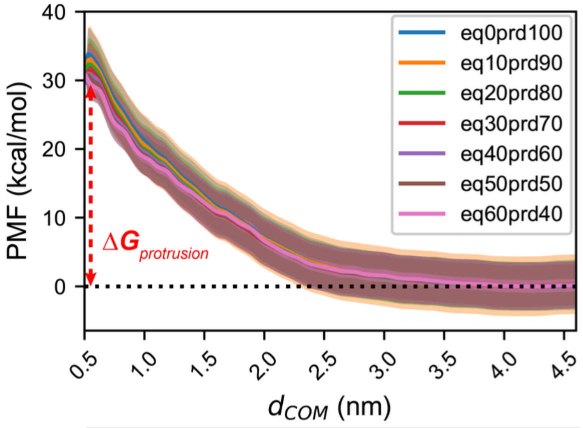


| Consecutive intervals | RMSE |
|---|---|
| eq0prd100 & eq10prd90 | 1.472 |
| eq10prd90 & eq20prd80 | 0.525 |
| eq20prd80 & eq30prd70 | 1.273 |
| eq30prd70 & **eq40prd60** | 0.185 |
| **eq40prd60** & eq50prd50 | 0.685 |
| eq50prd50 & eq60prd40 | 0.527 |

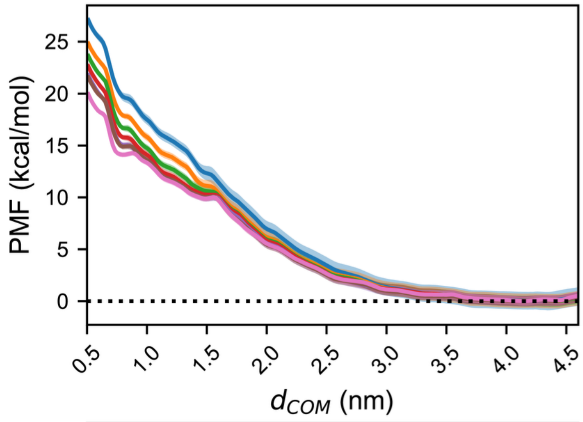


| Consecutive intervals | RMSE |
|---|---|
| eq0prd100 & eq10prd90 | 1.757 |
| eq10prd90 & eq20prd80 | 0.960 |
| eq20prd80 & eq30prd70 | 0.843 |
| eq30prd70 & **eq40prd60** | 0.670 |
| **eq40prd60** & eq50prd50 | 0.175 |
| eq50prd50 & eq60prd40 | 1.584 |

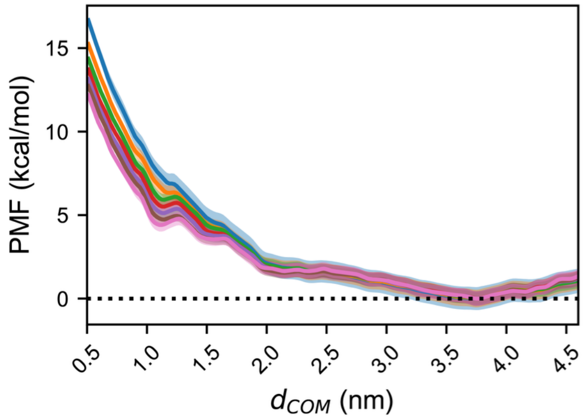


| Consecutive intervals | RMSE |
|---|---|
| eq0prd100 & eq10prd90 | 1.271 |
| eq10prd90 & eq20prd80 | 0.826 |
| eq20prd80 & eq30prd70 | 0.557 |
| eq30prd70 & **eq40prd60** | 0.480 |
| **eq40prd60** & eq50prd50 | 0.332 |
| eq50prd50 & eq60prd40 | 0.527 |

**Supplementary Figure S22.** Umbrella sampling (US) analysis of unsaturated PIP tail protrusion toward the GPx4 40s catalytic loop in DOPC + PIP membranes. **A.** Representative snapshots illustrating the protrusion of the native PIP unsaturated lipid tail toward the GPx4 40s loop. The color scheme follows the same definition used in **Supplementary Figure S18**. Red dashed arrows and corresponding annotations indicate the distance between the center-of-mass of the GPx4 40s loop and the unsaturated lipid terminus of PIP, $d_{COM}$. **B.** Convergence tests for the protrusion of the unsaturated PIP tail. For each replica, US was performed for 100 ns in total and the converged interval for each replica is indicated in bold in the table. These converged intervals were chosen for the final WHAM calculation. The resulting free energy values ($\Delta G_{protrusion}$) for the three replicas are 31.01 ± 3.65 kcal/mol, 21.79 ± 0.50 kcal/mol, and 13.15 ± 0.43 kcal/mol.

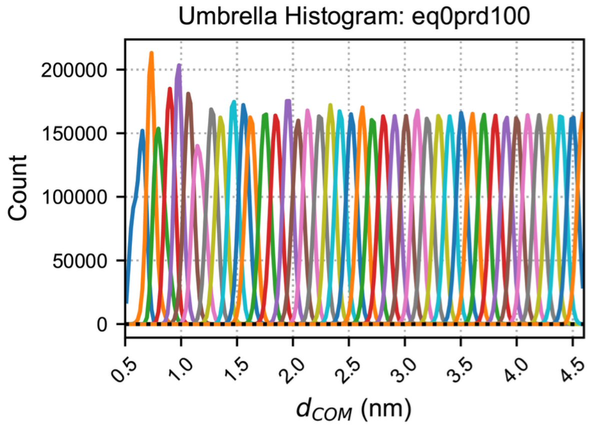
Umbrella Histogram: eq0prd100
Count
0
50000
100000
150000
200000
0.5 1.0 1.5 2.0 2.5 3.0 3.5 4.0 4.5
$d_{COM}$ (nm)

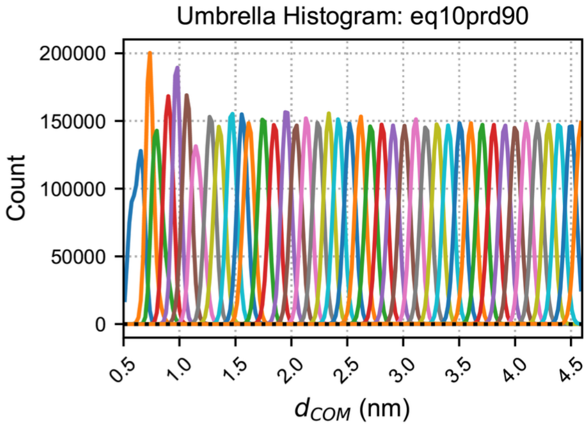
Umbrella Histogram: eq10prd90
Count
0
50000
100000
150000
200000
0.5 1.0 1.5 2.0 2.5 3.0 3.5 4.0 4.5
$d_{COM}$ (nm)

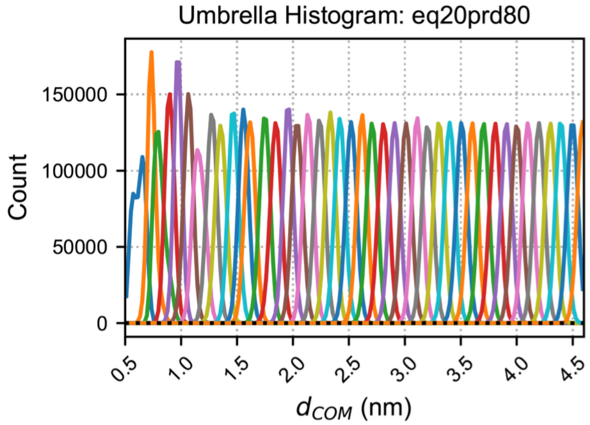
Umbrella Histogram: eq20prd80
Count
0
50000
100000
150000
0.5 1.0 1.5 2.0 2.5 3.0 3.5 4.0 4.5
$d_{COM}$ (nm)

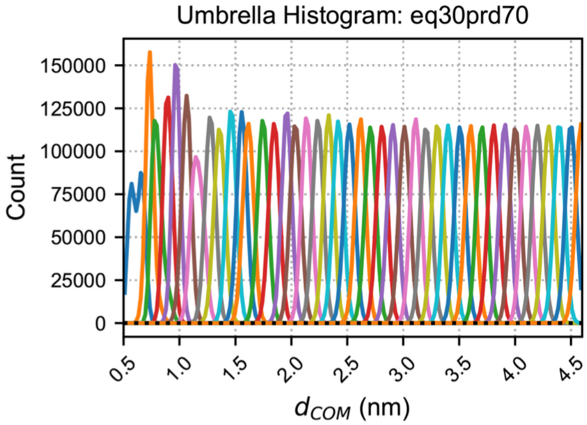
Umbrella Histogram: eq30prd70
Count
0
25000
50000
75000
100000
125000
150000
0.5 1.0 1.5 2.0 2.5 3.0 3.5 4.0 4.5
$d_{COM}$ (nm)

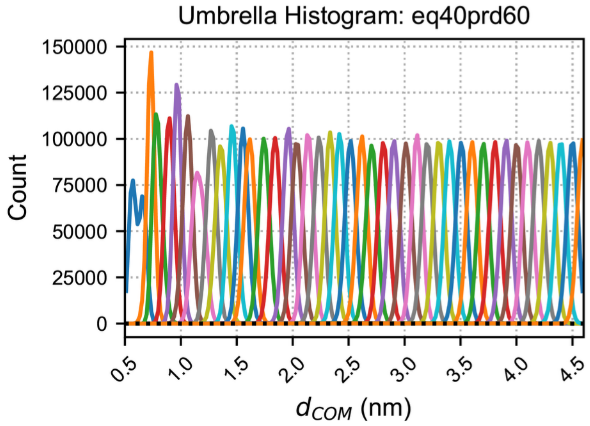
Umbrella Histogram: eq40prd60
Count
0
25000
50000
75000
100000
125000
150000
0.5 1.0 1.5 2.0 2.5 3.0 3.5 4.0 4.5
$d_{COM}$ (nm)

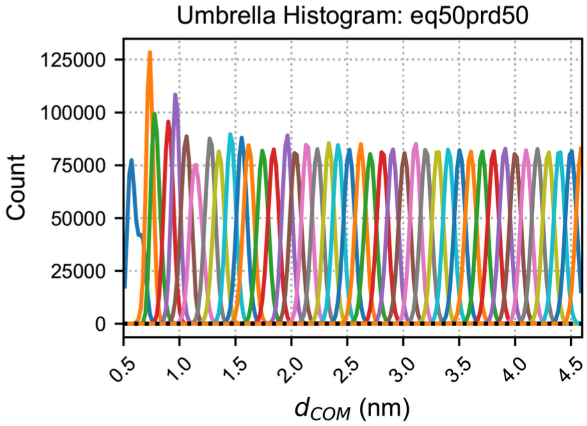
Umbrella Histogram: eq50prd50
Count
0
25000
50000
75000
100000
125000
0.5 1.0 1.5 2.0 2.5 3.0 3.5 4.0 4.5
$d_{COM}$ (nm)

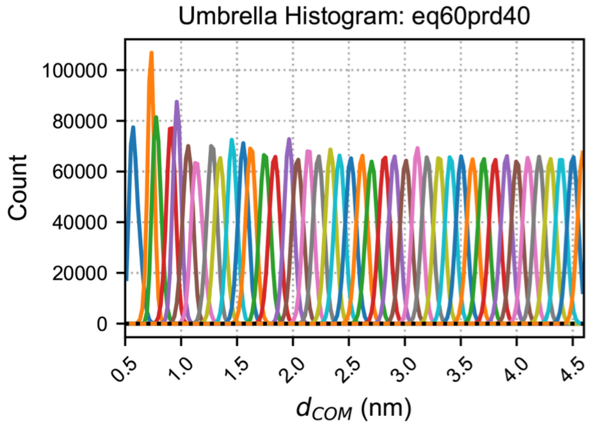
Umbrella Histogram: eq60prd40
Count
0
20000
40000
60000
80000
100000
0.5 1.0 1.5 2.0 2.5 3.0 3.5 4.0 4.5
$d_{COM}$ (nm)

**Supplementary Figure S23.** Histogram overlap across umbrella sampling windows for PIP tail protrusion. Data represent the unsaturated PIP tail protrusion toward the GPx4 40s loop in DOPC + PIP membrane systems. Histograms are shown for successive equilibration and production time intervals to demonstrate sampling overlap. Results for Replica 1 are shown as a representative example.

**A.**

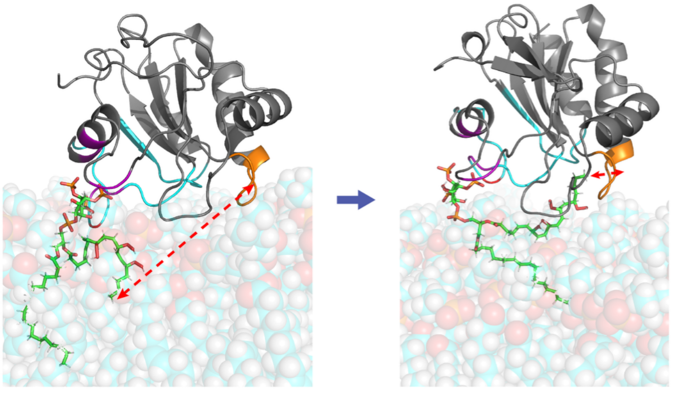


**B.**

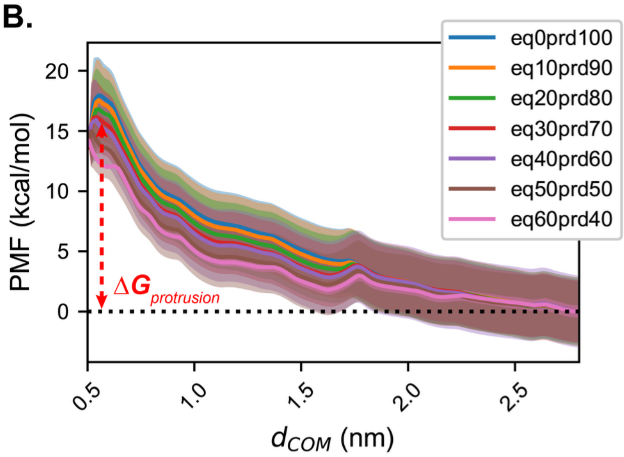


| Consecutive intervals | RMSE |
|---|---|
| eq0prd100 & **eq10prd90** | 0.196 |
| **eq10prd90** & eq20prd80 | 0.313 |
| eq20prd80 & eq30prd70 | 0.322 |
| eq30prd70 & eq40prd60 | 0.645 |
| eq40prd60 & eq50prd50 | 0.647 |
| eq50prd50 & eq60prd40 | 0.577 |

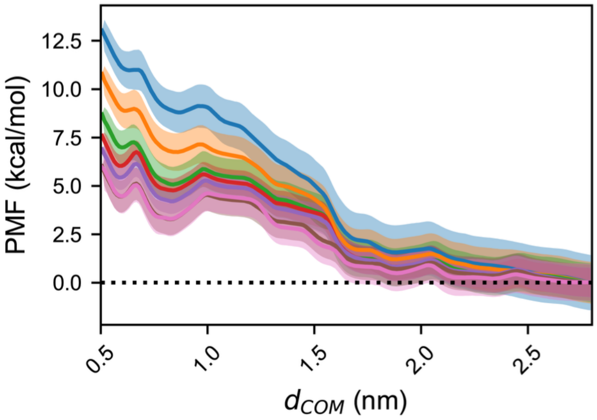


| Consecutive intervals | RMSE |
|---|---|
| eq0prd100 & eq10prd90 | 1.566 |
| eq10prd90 & eq20prd80 | 1.488 |
| eq20prd80 & eq30prd70 | 0.953 |
| eq30prd70 & eq40prd60 | 0.596 |
| eq40prd60 & **eq50prd50** | 0.567 |
| **eq50prd50** & eq60prd40 | 0.140 |

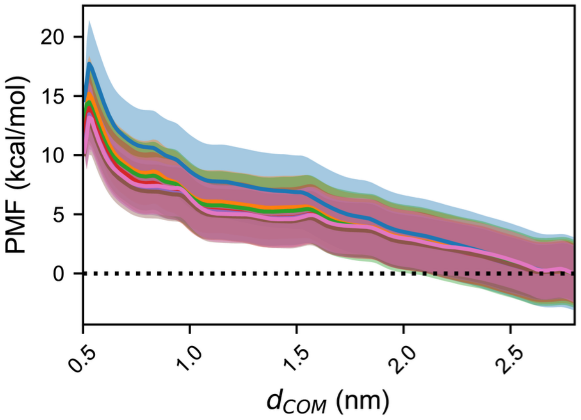


| Consecutive intervals | RMSE |
|---|---|
| eq0prd100 & eq10prd90 | 1.404 |
| eq10prd90 & eq20prd80 | 0.411 |
| eq20prd80 & eq30prd70 | 0.803 |
| eq30prd70 & eq40prd60 | 0.916 |
| eq40prd60 & **eq50prd50** | 0.289 |
| **eq50prd50** & eq60prd40 | 0.148 |

**Supplementary Figure S24.** Umbrella sampling (US) analysis of unsaturated PIP tail protrusion toward GPx4 40s catalytic loop in DOPC + PIP hydroperoxide membrane systems. **A.** Representative snapshots illustrating the protrusion of the peroxidized PIP unsaturated lipid tail toward the GPx4 40s loop. The color scheme follows the same definition in **Supplementary Figure S18**. Red dashed arrows and corresponding annotations indicate $d_{COM}$. **B.** Convergence tests for the protrusion of the unsaturated PIP tail. For each replica, US was performed for 100 ns in total and the converged interval for each replica is indicated in bold in the table. These converged intervals were chosen for the final WHAM calculation. The resulting free energy values ($\Delta G_{protrusion}$) for the three replicas are 17.56 ± 3.40 kcal/mol, 6.02 ± 0.83 kcal/mol, and 12.81 ± 2.77 kcal/mol.

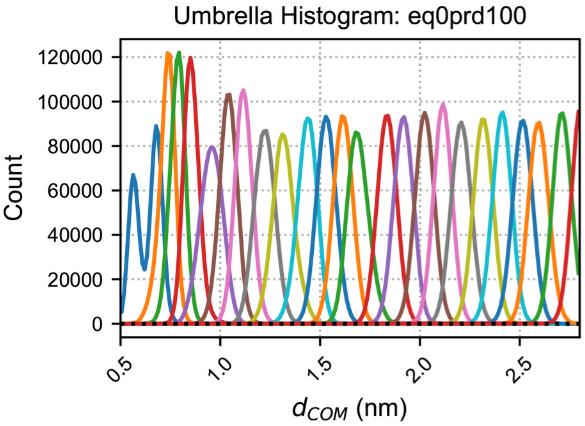
Umbrella Histogram: eq0prd100
Count
$d_{COM}$ (nm)

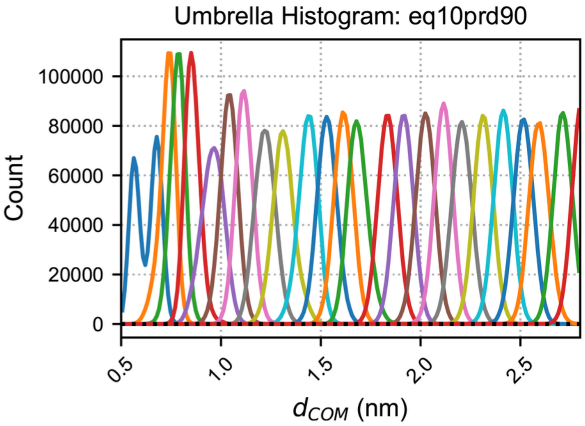
Umbrella Histogram: eq10prd90
Count
$d_{COM}$ (nm)

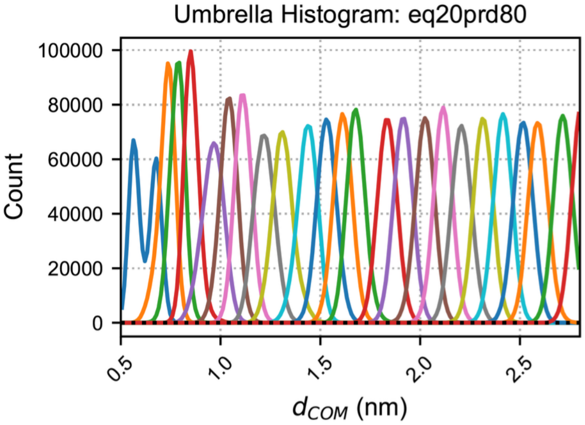
Umbrella Histogram: eq20prd80
Count
$d_{COM}$ (nm)

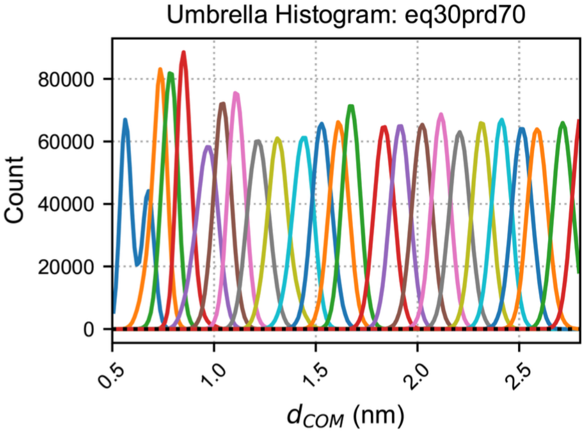
Umbrella Histogram: eq30prd70
Count
$d_{COM}$ (nm)

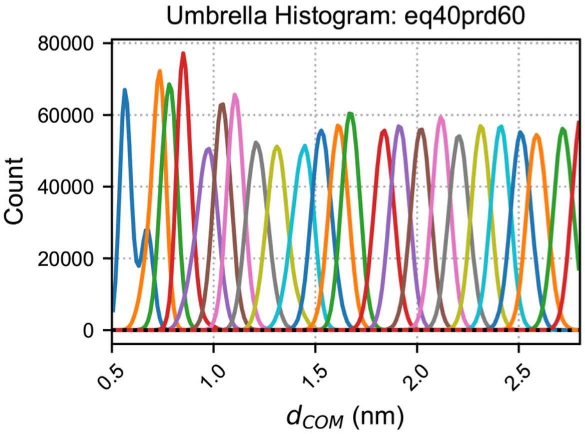
Umbrella Histogram: eq40prd60
Count
$d_{COM}$ (nm)

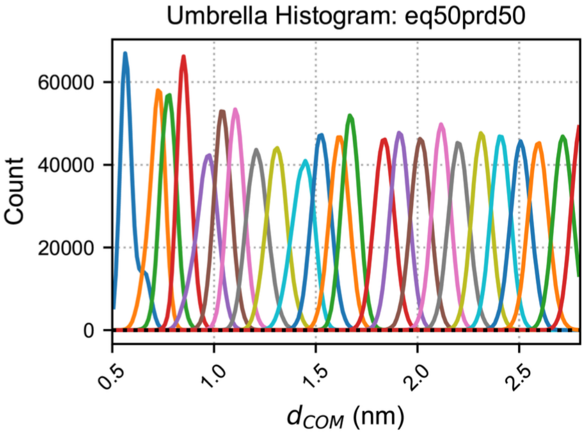
Umbrella Histogram: eq50prd50
Count
$d_{COM}$ (nm)

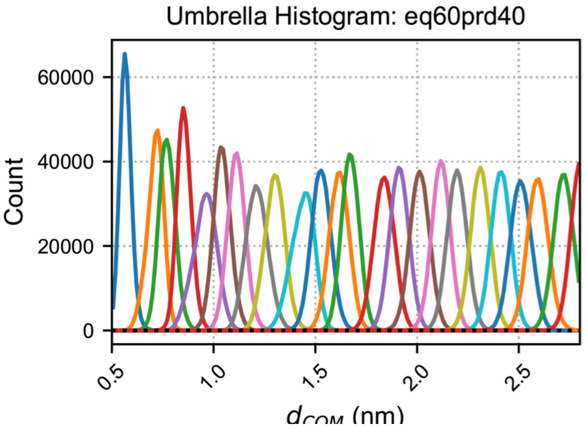
Umbrella Histogram: eq60prd40
Count
$d_{COM}$ (nm)

**Supplementary Figure S25.** Histogram overlap across umbrella sampling windows for peroxidized PIP tail protrusion. Data represent the peroxidized unsaturated PIP tail protrusion toward the GPx4 40s loop in DOPC + PIP-hydroperoxide membrane systems. Histograms are shown for successive equilibration and production time intervals to demonstrate sufficient window overlap. Results for Replica 1 are shown as a representative example.
